# Pattern-triggered immunity in blue and white seed cultivars of *Papaver somniferum*

**DOI:** 10.1101/2025.02.23.639761

**Authors:** Jhonny Stalyn Hernandéz Orozco, Oksana Iakovenko, Adam Zeiner, Marie Hronková, Jiří Kubásek, Bára Kučerová, Iveta Vachová, Serban Pop, Petr Maršík, Markéta Macho, Pavla Fojtíková, Andrea Rychlá, Ondřej Hejna, Ivan Kulich, Michael Wrzaczek, Martin Janda

## Abstract

*Papaver somniferum* (poppy) is a traditional component of Central and Eastern European cuisine and an important oilseed crop in the region. The crucial thread for poppy stable yield is pathogen infection. Thus, we need to understand poppy defence mechanisms in detail. The first robust layer of plant immunity, which plays a crucial role in combat against pathogens, is pattern-triggered immunity (PTI). Here, we provide the first insights into PTI in poppy. We selected four poppy varieties used in the food industry. We investigated poppy response to various peptide elicitors acting as microbe-associated molecular patterns (MAMPs) and damage-associated molecular patterns (DAMPs). Flg22 induced the most robust reactive oxygen species (ROS) burst among all tested peptides. Flg22 also triggered putative mitogen-activated protein kinase (MAPK) phosphorylation and seedling growth inhibition in all tested cultivars. We identified *PsWRKY22* and *PsPR2* as candidate marker genes suitable for monitoring poppy PTI responses. The tested poppy cultivars have low levels of salicylic acid. Callose accumulation was triggered by wounding but not by flg22. For studying PTI in poppy, wounding is a challenge that needs to be considered as it can obscure potential PTI responses. Our findings highlight conserved aspects of poppy immunity and challenges in studying poppy PTI. The established pipeline facilitates improving our understanding of poppy immunity and has the potential for widespread application in poppy breeding and improving selection for broad-spectrum disease resistance provided by enhanced PTI.

**GRAPHICAL ABSTRACT:** 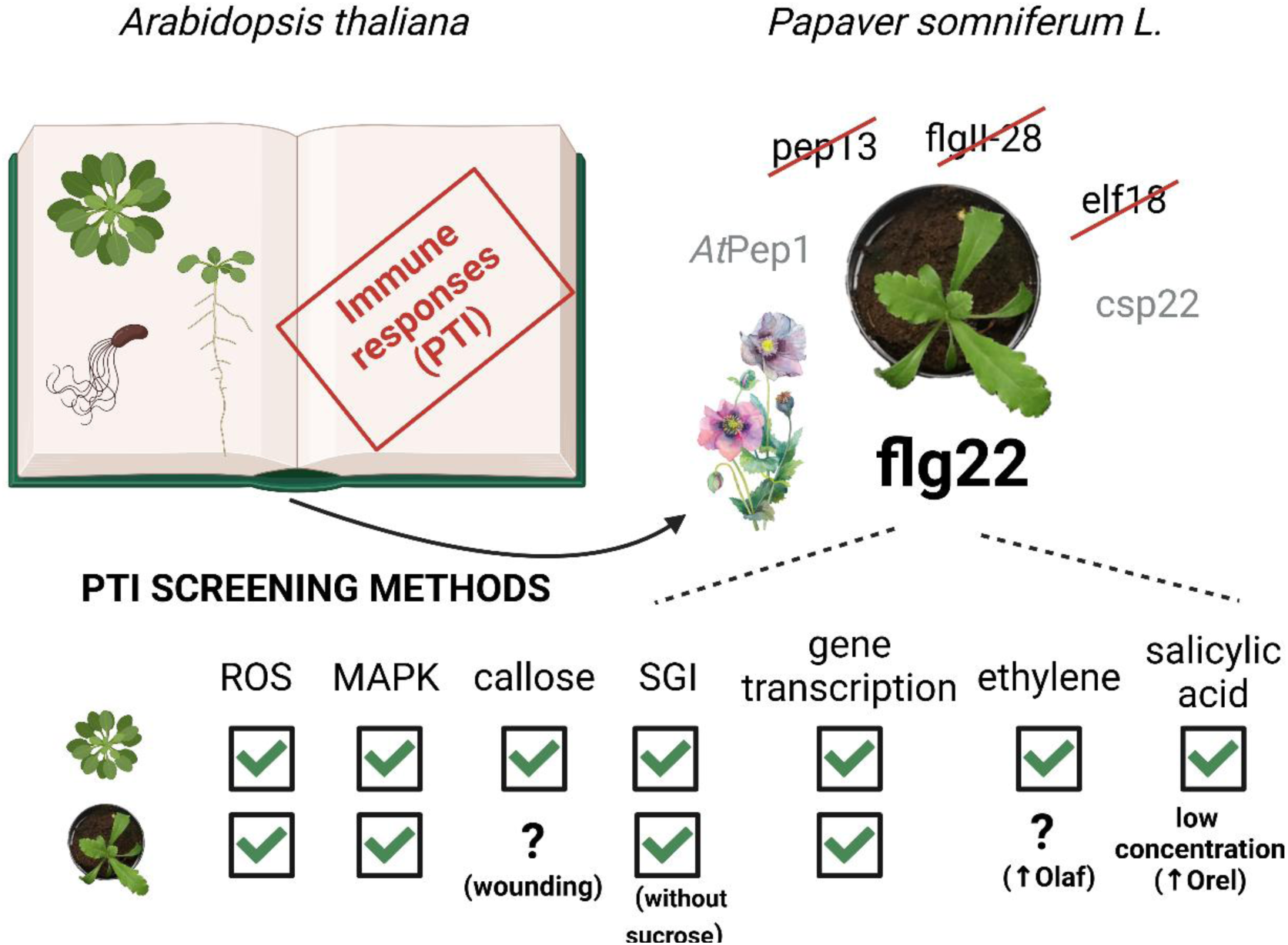
Graphical abstract. The establishment of the methods for studying pattern-triggered immunity (PTI) in *Papaver somniferum* L. (poppy) was inspired by the knowledge of the model plant *Arabidopsis thaliana*. The study showed a similarity between Arabidopsis and poppy in response to flg22 but also pointed out the obstacles for PTI analysis in poppy and the differences compared to the model plant. *Created with BioRender.com*.

## Introduction

Poppy cultivation has a long-standing history, with records dating back to 4000 BC when the ancient Sumerians cultivated it. Poppy was called *Hul Gil* ("flower of joy") (museum.dea.gov, 2025). Due to its high content of alkaloids, with more than 80 identified in *Papaver somniferum* (www.efsa.europa.eu, 2025). Today, poppy remains important as both a medicinal and pharmaceutical crop (Beaudoin & Facchini, 2014; Singh *et al*., 2019). In Central and Eastern Europe, it is a significant oilseed crop (Neugschwandtner *et al*., 2023). The oil content of poppy seeds is around 40-50 %, rich in vitamin E and minerals (Melo *et al*., 2022). The Czech Republic is the world’s leading producer and exporter of poppy seeds per capita, contributing $88 million in exports in 2007 (Prochazka & Smutka, 2012). Based on the Czech Statistical Office, in 2024, breadseed poppy covered 36 611 hectares in the Czech Republic, which is almost double compared to the potatoes area (22 747 hectares), and the export in 2021 was around 60 million € (csu.gov.cz, 2025).

Based on United Nations estimation, globally, up to 40 % of total crop production is lost to pests annually, with diseases costing an estimated $220 billion and invasive insects adding another $70 billion (www.fao.org). Like other crops, poppy is vulnerable to pathogens and pests (Bailey *et al*., 2000; Thangavel *et al*., 2018). Breadseed poppy, bred for its low alkaloid content, has a reduced natural defence capabilities. Significant diseases include leaf blight, caused by the fungal pathogen *Pleospora papaveracea* (O’Neill *et al*., 2000), and poppy downy mildew caused by the oomycete *Peronospora arborescens* (Landa *et al*., 2007), both of which can devastate entire fields. Bacterial diseases such as bacterial blight, caused by *Xanthomonas campestris* pv. *Papavericola* (Gingeras *et al*., 1978) and bacterial stem rot, caused by *Erwinia carotovora* (Alam *et al*., 2014), also threaten poppy plants.

Understanding plant immunity is crucial for mitigating the losses caused by biotic stress in sustainable agriculture. In the past thirty years, research on plant immunity, particularly in model plants like *Arabidopsis thaliana*, has provided invaluable insights. A key aspect of plant immunity is the recognition of pathogens. Plants detect two main types of pathogen molecules: (i) pathogen-associated molecular patterns (PAMPs) and (ii) effectors. PAMPs are recognised by pattern recognition receptors (PRRs), mainly located on the plasma membrane, while effectors are typically detected in the cytosol by receptor-like kinases (RLK) (Jones & Dangl, 2006). PAMP recognition triggers the first layer of plant immunity, known as pattern-triggered immunity (PTI), which is the subject of this study in poppy. PAMPs are conserved molecules essential to pathogens and are chemically distinct from host molecules, ranging from peptides (e.g., flagellin, elf18) and sugars (e.g., chitin) to short-chain fatty acids (Bigeard *et al*., 2015; Boutrot & Zipfel, 2017). PTI can also be triggered by damage-associated molecular patterns (DAMPs) derived from plant molecules like extracellular ATP, cell wall fragments or peptides produced by plants under stress (Tanaka & Heil, 2021).

One of the best-studied ligand-receptor pairs involved in PTI is flg22, a 22-amino acid epitope of bacterial flagellin, and its receptor FLAGELLIN SENSING 2 (FLS2) (Tena, 2019). Discovered in 1999 (Felix *et al*., 1999; Gómez-Gómez *et al*., 1999). Research in *Arabidopsis thaliana* has revealed the molecular mechanism of flg22 binding to FLS2, including the involvement of the co-receptor BAK1 (Sun *et al*., 2013). This interaction triggers several typical defence responses, such as transient Ca^2+^ spikes (Chi *et al*., 2021), apoplast alkalisation (Felix et al., 1999), reactive oxygen species (ROS) bursts (Smith & Heese, 2014), mitogen-activated protein kinases (MAPK) cascade activation (Tian *et al*., 2021), an increase of callose deposition (Ellinger & Voigt, 2014), and changes in phytohormone levels, including increases in salicylic acid (Tsuda *et al*., 2008) and ethylene production (Felix *et al*., 1999), as well as dynamic transcriptional changes (Zipfel *et al*., 2004; Bjornson *et al*., 2021). Responses to flg22 have been observed in numerous plant species, including members of the *Solanaceae* and *Brassicaceae* families (Nguyen *et al*., 2010; Lloyd *et al*., 2014), as well as in rice (Takai *et al*., 2008). The detailed understanding of PTI has demonstrated a significant potential for plant breeding and biotechnology. For example, the transfer of the PRR EF-Tu receptor (EFR) from *Arabidopsis*, which recognises a peptide from the bacterial elongation factor elf18 to tomato has significantly increased resistance to bacterial pathogen *Ralstonia solanaceae* (Lacombe *et al*., 2010). To our best knowledge, no systematic study focusing on PAMPs or DAMPs, including flg22, has so far been conducted on breadseed poppy (*Papaver somniferum*).

This study aims to establish, optimise, and present a methodology suitable for the analysis of the PTI responses in poppy. We observed that flg22 is the most potent of the tested peptide PAMPs or DAMP. Thus, we used flg22 as a representative PTI elicitor. Our results represent the first picture of poppy immune responses and form a basis for PTI research in poppy, with possible usage in poppy breeding efforts for increased resistance to pathogens, in future.

## MATERIALS AND METHODS

### Plant material

In this study were used four *Papaver somniferum L.* cultivars provided by OSEVA PRO s.r.o (Opava, Czech Republic): Gerlach (ID nr. 15O0800148), Orel (ID nr. 15O0800187), Opex (ID nr. 15O0800203), and Olaf (new cultivar, in time of manuscript preparation without assigned ID number). Plants were grown in two distinct styles for the experiments: *in vitro* (sterile conditions; seedlings) and in soil (non-sterile conditions; adult plants). For both cultivation styles, seedlings were sterilised using 30% (v/v) sodium hypochlorite solution for 6-10 minutes and four times washed using sterile distilled water (*in vitro* conditions) or distilled water (in soil conditions). Sterilised seeds were stratified (4 °C) in dark in water for 1-5 days prior sowing. The similar growing conditions as for poppy were used for *Arabidopsis thaliana* plants. Potato plants were grown in soil in the greenhouse under not controlled light conditions, and only maximum temperature was controlled under 26 °C. 3-6 weeks old potato plants were used for the experiments. Potatoes were used as a control and not for main results.

*<u>In soil</u>* <u>conditions</u>, seeds were sowed into hydroponia in perlite and watered with hydroponic nutrient solution containing Jungle Garden Base 0.1% (v/v) and Jungle Garden G1 0.25% (v/v) (JUNGLE Indabox, Czech Republic). After 6-9 days, seedlings were transferred into the peat pellet (Jiffy 7, Bohemiaseed, Czech Republic) (Fig. S1). The plants were watered with distilled water approximately once a week. Plants were grown for 5 - 6 weeks in Phytotron (Photon Systems Instruments, Czech Republic) under a short-day photoperiod (10 h day/14 h day/night regime), light intensity of 120 µmol·m^-2^·s^-1^, temperature 22 °C /18 °C day/night and humidity ∼60%.

*<u>In vitro</u>* <u>conditions</u>, seeds were sown on solid Murashige and Skoog medium (MS), including vitamins (Duchefa, Netherlands) containing 1% (w/v) sucrose and 1.2% (w/v) Gelrite (Duchefa, Netherlands), pH was adjusted to 5.6-5-7 (Fig. S2). The seedlings grew on solid media for 2-5 days under 16 h/8 h day/night regime, light intensity 110 - 140 µmol·m^-2^·s^-1^ and temperature 23-25 °C. Then, seedlings were moved to the 24-well plate with 400 µL of MS liquid media, including vitamins and 1% (w/v) sucrose, and grew under the same growing conditions as on solid media. In vitro seedlings were used for the ROS analysis, seedlings growth inhibition assay (for this assay was used also liquid MS media without sucrose) and callose measurement (further details in dedicated sections).

### Measurement of ROS production using luminol-based method

The ROS production was determined using the luminol-based assay as previously described (Janda *et al*., 2019). Briefly,

<u>ROS measurement in leaves from adult poppy plants grown on soil (in soil conditions</u>): The leaf discs (4 mm in diameter) of 5-6-week-old poppy plants were incubated in 100 µL of distilled water into the white 96-well plate and in the dark in room temperature overnight. The water was replaced with the solution containing 200 µM luminol (Serva, 28085.02), 20 µg/mL horseradish peroxidase (HRP, Apollo Scientific, Aposbitp1327) and particular elicitor (concentration of used elicitors is specified in results section and figure legends): flg22 (EZBiolab; QRLSTGSRINSAKDDAAGLQIA; (Chinchilla *et al*., 2006)), *Xcc*Flg22 (Nzytech; QRLSSGLRINSAKDDAAGLAIS; this study), flgII-28 (Chinese Peptide; ESTNILQRMRELAVQSRNDSNSATDREA; (Moroz & Tanaka, 2020)), elf18 (EZBiolab; Ac-SKEKFERTKPHVNVGTIG; (Kunze *et al*., 2004)), *At*Pep1 (Chinese Peptide; ATKVKAKQRGKEKVSSGRPGQHN; (Ortiz-Morea *et al*., 2016)), csp22 (Chinese Peptide; AVGTVKWFNAEKGFGFITPDDG; (Trinh *et al*., 2024)) and Pep13 (Chinese Peptide; VWNQPVRGFKVYE.; (Nietzschmann *et al*., 2019)) to induce the production of ROS, which is monitored as luminescence intensity correlating with the concentration of H_2_O_2_ and described as relative luminescence unit (RLU) or as the area under the curve from the 4^th^-24^th^ minute (total photon counts). For luminescence measurement, a luminometer Tecan SparkCyto 300 (Tecan, Switzerland) was used.

<u>ROS measurement in seedlings grew *in vitro*</u>: Five-days old seedlings grown on the solid medium in sterile conditions were moved to MS liquid medium (described above) and put into 24-well plates for 24 hours. Then, rinsed individual seedlings were moved into a white bottom 96 well plate with 200 µL distilled water and kept in the dark for at least 16 h. After at least 16 hours in the dark water was replaced by 200 µL of 5 µM flg22, HRP (20 µg/mL), luminol (200 µM), and distilled water. Seedlings treated with distilled water instead of flg22 were used as control.

### Measurement of ROS in poppy root apoplast using Amplex Red

The ROS production in roots was determined using Amplex Red Reagent (Invitrogen, A12222) method as previously described in (Kulich *et al*., 2025). The FV1000 Olympus Confocal Microscope with objective UPlanSApo 10x/0.40 (wavelengths: excitation 559 nm; emission 583 nm) was used for imaging. For experiment were used roots of three to four days old poppy seedlings growing on 1/2 MS media including vitamins, 1% (w/v) plant agar and 1% (w/v) sucrose. Poppy seedlings were placed on the microscopic glass slide in a drop of 1/2 MS media and were left to rest for approximately 2 minutes, followed by the first was by mock medium wash. After this, a second medium containing 1 µM flg22 was added, replacing the original media. Media replacement was observed as a rapid drop in the intensity of the Amplex Red signal roughly to the levels of the control Amplex Red sample. This stain predominantly localises to the apoplast close to elongation zone and exhibits red fluorescence halo upon oxidation to resorufin in the presence of H_2_O_2_. Images were captured every 3 s. Fiji ImageJ (version 8) was used for the image analysis. Intensity of stain on ROI localised close to the apoplast of elongation zone were measured. Mean grey value at the time 0 s is subtracted from all following values. The data was not normalised.

### MAPK activation assay

For analysis of the putative MAPK phosphorylation in poppy we used leaves from 5-6 weeks old plants. We used two styles of 5µM flg22 treatments:

<u>Leaf infiltration with needleless syringae</u>. Non-treated leaves, parts of leaves infiltrated with distilled H_2_O or parts of leaves infiltrated with 5µM flg22 were collected 15 and 30 minutes after the infiltration frozen in liquid nitrogen.

<u>Leaf discs treatment</u>. Leaf discs (4 mm) were either immediately frozen in liquid nitrogen (steady negative control), or incubated at least 16 h in distilled water under continuous dark and treated by changing of distilled H_2_O either with fresh distilled H_2_O or with water solution containing 5µM flg22. The treatment was performed for 15 and 30 minutes. After that the leaf discs were frozen in liquid nitrogen.

Collected samples were homogenised using mortar and pestle in liquid nitrogen. Proteins were extracted (50 mM HEPES, pH 7.5; 75 mM NaCl; 1 mM EGTA; 1 mM MgCl_2_; 1 mM NaF; 10% (v/v) glycerol; 1mM DTT; cOmplete, EDTA-free Protease Inhibitor Cocktail (Roche, 11873580001) and Pierce Phosphatase Inhibitor Mini Tablets (Thermo Scientific, A32957)) and quantified with Bradford assay (Bradford, 1976). 15 µg of total protein was separated by 12% SDS-PAGE and transferred to PVDF membrane (Immun-Blot Low Fluorescence PVDF Membrane, BioRad, 1620264). Membrane was blocked with 5% (w/v) bovine serum albumin (BSA) in Tris-buffered-salin (TBS)-0.1% (v/v) Tween-20 (T) (1 hour at room temperature, or overnight at 4 °C), incubated with primary antibody (Phospho-p44/42 MAPK (Erk1/2) (Thr202/Tyr204) Antibody, Cell Signaling Technology, #9101; diluted 1:1000 in 1% (w/v) BSA in TBS-T, overnight at 4 °C), washed in TBS-T (5 times 5 minutes at room temperature), incubated with secondary antibody (StarBright™ Blue 520 Goat Anti-Rabbit IgG, BioRad, #12005870; diluted 1:2500 in 1% (w/v) BSA in TBS-T, 30 minutes at room temperature), and finally washed in TBS (5 times 5 minutes at room temperature). The fluorescent antibody was detected with documentation unit ChemiDoc (BioRad) with the provided protocol for the detection of StarBright Blue 520. The loading control is represented as Coomassie dye (0.25% (w/v) Coomassie Brilliant Blue R-250, 45% (v/v) methanol, 10% (v/v) glacial acetic acid) stained, and destained (45% (v/v) methanol, 10% (v/v) glacial acetic acid) membrane, which was documented with already declared documentation unit. Analysis was repeated 2 times with similar result. Final images were analysed with Image Lab (BioRad, 6.0.1).

### Seedlings growth inhibition assay

Two to five-days old poppy (or Arabidopsis) seedlings grew in full MS including vitamins solid medium containing 1% sucrose (*in vitro* conditions) were moved to 24-well plates containing 500 µL MS liquid medium including vitamins with or without 1% (w/v) sucrose (Duchefa, Netherlands) either supplemented with 5 µM flg22 or sterile H_2_O. One to three seedlings were placed into each well. Medium +/- flg22 was refreshed after 3 days with MS medium with sucrose. In experiments with MS without sucrose, we did not refresh the media. Seedlings were cultivated for 5-7 days. Thus, 10-12 days old seedlings were harvested for analysis. The seedlings were weighed individually.

### Gene expression analysis

For analysis of the gene expression in poppy we used leaves from 5-6 weeks old plants and leaf discs treatment (similar to ROS assay method) was performed with distilled H_2_O (mock) or with 5 µM flg22 for indicated time. Leaf discs were either immediately frozen (steady negative control), or incubated overnight in distilled water and treated with fresh distilled H_2_O or with water solution containing 5µM flg22 for corresponding time. After that the leaf discs were frozen in liquid nitrogen. Three to four biological replicates for each treatment and each time point were prepared.

Samples were homogenised in tubes with 1.3 mm silica beads using a FastPrep-24 instrument (MP Biomedicals, USA). Total RNA was extracted from leaves or discs of P. somniferum using FavorPrep Total RNA Isolation Kit (FAVORGEN Biotech Corp.) according to the produceŕs instructions. Part of RNA samples was treated with DNA free Kit, DNAse Treatment & Removal (Invitrogen, AM1906) to eliminate genomic DNA contamination, and 1 µg of pure total RNA was used for the synthesis of cDNA by High Capacity cDNA Reverse Transcription Kit (Applied Biosystems, 4368814) according to manufactureŕs instructions. RT-qPCR was performed with an qTower3 Real-time PCR detection system (Analytic Jena) using HOT FIREPol Eva Green qPCR Mix Plus ROX (SOLIS BIODYNE). 4 µL (1:20 diluted) of the RT product in a final reaction volume of 20 µL was used. The following PCR program was used throughout the study: 95°C for 12 min, followed by 40 cycles of 95°C for 15 s, 55°C for 20 s and 72°C for 15 s.Two technical replicates were set up for each cDNA template. Data were normalised to the reference gene Actin and to the transcript level relative to the non-treated leaves or discs for each sample by the comparative CT method (Livak & Schmittgen, 2001). The primers used in qRT-PCR are listed in Table S1.

### Callose accumulation analysis

Treatment by flg22 for callose accumulation analysis in poppy plants was done in three ways: using 8-day-old seedlings grown *in vitro*, leaf discs and whole fully developed leaves from 4-5 weeks old plants. Seedlings growing in the same way as for seedlings growth inhibition assay were transferred for 72 h to fresh MS medium including vitamins and with 1% (w/v) sucrose for mock and same medium containing 5µM flg22. For the leaf infiltration, three leaves at the same developmental stage from 4-6 plants were infiltrated with 5 µM flg22 solution, mock was distillate water. After 72 h treated, mock leaves and three leaves per plant from four no-treated plants were detached for further staining procedure. In the leaf disc treatment, 4 mm discs were cut from three leaves of four poppy plants and placed in a 96-well plate, one disk to one well with 0.1 mL of distillate water. After overnight incubation in the dark room temperature, the solution was replaced by 5 µM flg22 or fresh distilled water for mock for 12 hours. No treated discs were cut immediately before staining.

After treatment, plant material was fixed in 96% ethanol: glacial acetic acid (3:1, v/v), the solution was replaced several times until plant tissue was decolorated. After that, it was rehydrated and sequentially incubated in 70/ 50/ 30 % (v/v) ethanol and distilled water for 1 hour in each solution. Finally, fixed tissue was stained with 0.01 % (w/v) aniline blue in 150 mM K_2_HPO_4_, pH 9.5 overnight.

Callose deposition was observed using DAPI channel (wavelengths: excitation 359 nm/emission 455 nm), BX63 OlympusWidefield Fluorescence Microscope, 4X objective Olympus UPlanXApo 4X/0.16. The ratio of all callose spots area on the whole leaf or disc area was calculated using Labkit in Fiji ImageJ 8 software (Schindelin *et al*., 2012).

### Poppy FLS2 receptor and flg22 binding analysis

FLS2 homolog protein of *P. somniferum L.* was identified using as query *A. thaliana* FLS2 protein sequence (AT5G46330.1) through Blastp (Altschul *et al*., 1990). Blastp was set up to search for FLS2 only in *Papaver somniferum L*. (NCBI ID ASM357369v1), the hit protein named as LRR receptor-like serine/threonine-protein kinase FLS2 (NCBI ID XP_026454075.1) was selected for modelling.

The amino acid sequence of putative *Ps*FLS2 was used for *in silico* 3D modelling using AlphaFold2 server version 2.3.0 (Jumper *et al*., 2021). In the *Ps*FLS2 modelled structure, only the extracellular domain (LRRs region) was conserved.

*Xcc*flg22 peptide sequence was obtained through Blastp using as query sequence *Pseudomonas syringae* pv*. tomato str. DC3000* flagellin (NCBI ID AAO55467.1). Hit protein sequence in *Xanthomonas campestris* pv. *Campestris* named as flagellin (NCBI ID MEB1151700.1) was used. To obtain *Xc*flg22 3D structure the flg22 peptide structure from Sun et al. (2013) (RCSB PDB ID 4MN8) was used as a template and modelling was performed in PyMOL (Molecular Graphics System, Version 3.1 Schrödinger, LLC.).

*Ps*FLS2 and *Xcc*flg22 PDB files were treated using OpenBabelGUI 3.1.1 (O’Boyle *et al*., 2011) to add charges in the residues and convert PDB to PBDQT files. The PBDQT files were used for the molecular docking process through PyRx-Python Prescription 0.8 Virtual Screening Tool (Dallakyan & Olson, 2015) a visual interface of AutoDock4 v4.2.6 program (Morris *et al*., 2009). The autodock wizard was set up for docking, with a modified grid to cover the entire FLS2 section where the interaction with flg22 could occur. Grid area was determined by an alignment between the *Ps*FLS2 (XP_026454075.1) and *At*FLS2 (AT5G46330.1) protein sequences using MEGA11: Molecular Evolutionary Genetics Analysis version 11.0. 10 (Tamura *et al*., 2021), the alignment was performed with the MUSCLE algorithm (with default setting) (figure S7). Potential *Ps*FLS2 residues that could interact with *Xcc*flg22 or flg22 were selected using the list of residues in *At*FLS2 that interact with flg22 reported by (Sun *et al*., 2013).

In *At*FLS2 docking, 3D structure (RCSB PDB accession number 4MNA) was prepared by removing water and other molecules using PyMOL, and the resulting PBD file was prepared in the same manner as *Ps*FLS2. After the docking process, the ligand with the lowest binding energy (B.A.) was selected and for visualization and imaging of the complexes, open-source web-based toolkit Open Mol*Viewer (Sehnal *et al*., 2021) was used. B.A. was used to calculate *in silico* Kd (equilibrium dissociation constant) using the Gibbs-Helmholtz equation. .

### Salicylic acid measurement

We analysed salicylic acid (SA) concentration similarly as we did for Arabidopsis samples in our previous study (Leontovyčová *et al*., 2019; Pluhařová *et al*., 2019). Briefly, SA analysis was carried out in four biologically independent samples from each variant (treatment, cultivar), with every sample containing 100-200 mg fresh weight leaves from five to six-week-old poppy plants. Material from at least three plants was collected and considered one sample. Samples were homogenised in tubes with 1.3 mm silica beads using a FastPrep-24 instrument (MP Biomedicals, USA). The samples were then extracted with a methanol/H_2_O/formic acid (15:4:1, v:v:v) mixture supplemented with stable-isotope-labelled ^13^C-SA internal standards. Extracts were subjected to solid phase extraction using Oasis MCX cartridges (Waters Co., Milford, MA, United States) and eluted with methanol. The eluate was evaporated to dryness and dissolved in 15% (v/v) acetonitrile/water directly before the analysis. Quantification was performed on an Ultimate 3000 high-performance liquid chromatograph (UHPLC, Dionex; Thermo Fisher Scientific, Waltham, MA, United States) coupled to an IMPACT II Q-TOF ultra-high resolution and high-mass-accuracy mass spectrometer (HRAM-MS; Bruker Daltonik, Bremen, Germany). Separation was carried out using an Acclaim RSLC 120 C18 column (2.2 m, 2.1 × 100 mm; Thermo Fisher Scientific, Waltham, MA, United States) mobile phase consisting of 0.1% (v/v) formic acid in methanol by gradient elution. The full-scan data were recorded in negative electrospray ionization (ESI) mode.

### Ethylene measurement

The analysis of ethylene production was adapted from(Felix *et al*., 1999). In brief, six well-developed poppy or Arabidopsis leaves from 5-6 weeks old plants cultivated in soil were non-treated or infiltrated with distilled H_2_O or with 5µM flg22 and put into 20 mL glass vials containing 10 mL of distilled water. Vials were closed with silicone/PTFE septa and leaves were incubated at room temperature in the dark for four hours. Ethylene accumulated in the free air space was measured using gas chromatography. Ethylene was separated from atmospheric methane and other volatiles using PLOT column (Rt-Q-BOND, RESTEK, 30 m x 0.25 mm ID and 8.4 µm film thickness) under temperature of 38 °C and column flow 2 mL min^-1^ and detected by flame ionization detector (FID).

### Statistical data analysis and creation of graphs

For statistical data analysis and graph creation either MS Excel (Microsoft, 365 edition), or GraphPad Prism 10.0.0 (GraphPad Software, Boston, Massachusetts USA,) were used. Detail information about statistical analysis are available in particular figure legend. Figures from immunoblot Figure were prepared in Inkscape (Inkscape Developers, 1.3.2), but finalised in powerpoint.

## RESULTS

For our experiments, we selected four poppy cultivars: Gerlach (spring blue seed), Olaf (overwintering blue seed), Orel (spring white seed) and Opex (spring blue seed). Standard growing conditions were established using *in soil* peat pellets (Fig. S1) and *in vitro* sterile cultivation media (Fig. S2).

### ROS burst in *P. somniferum* leaves

ROS production is a rapid PTI response. We measured ROS using a luminol-based assay in leaf discs in 96-well plates (Smith & Heese, 2014). We treated poppy with five well-known peptide PAMPs (flg22, flgII-28, elf18, csp22, pep13) and one peptide DAMP (*At*Pep1). Among the peptides tested, flg22 from *Pseudomonas aeruginosa* (further referred to as flg22) triggered the most potent ROS burst in all cultivars (Fig.1A, B, C, D and S3), while *At*Pep1 and csp22 induced weak but detectable responses (Fig. 1B and S3). The peptides elf18, pep13, and flgII-28 did not elicit any observable ROS response (Fig. 1C and S3). For *At*Pep1, csp22, elf18, pep13 and flgII-28 we perform positive control using other plant species previously described in the literature (Fig. S4) (Moroz & Tanaka, 2020). Flg22 consistently triggered ROS production across all cultivars (Fig. 1D and S5). However, the response was more intensive in Arabidopsis compared to poppy, and a lower flg22 concentration was sufficient to saturate the response in Arabidopsis compared to poppy (Fig. 1E and S6).

**Figure 1.**
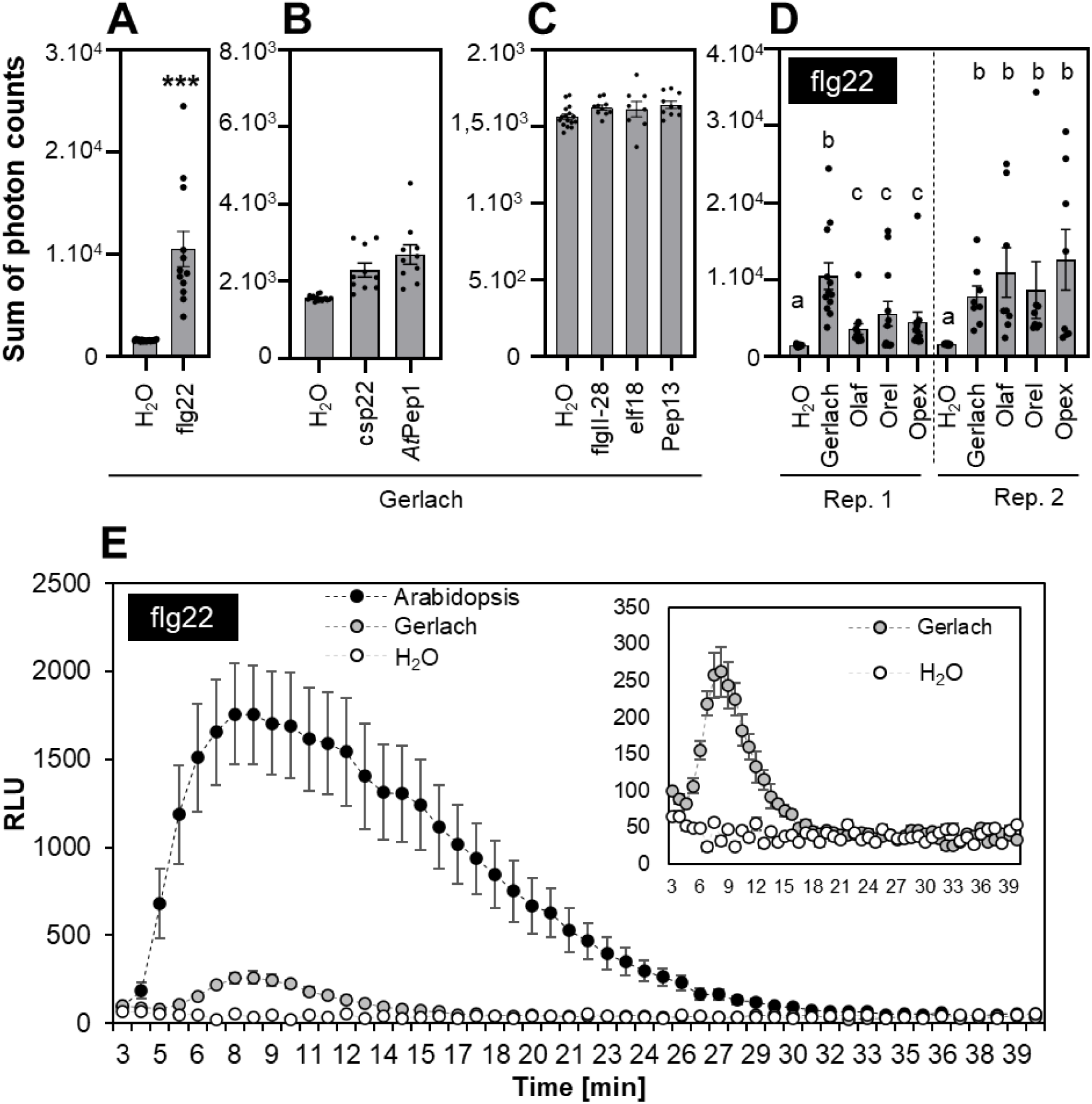
ROS burst in poppy after treatment with peptide elicitors. Discs were cut from 5-6 week-old poppy plants (A-E) or 5-6 week-old *A. thaliana* plants (E). **A-C)** Leaf discs from the Gerlach cultivar were treated with 1µM: flg22, flgII-28, AtuFlg22, csp22, *At*Pep1, elf18, and pep13. **D)** Discs from Gerlach, Olaf, Orel and Opex cultivars were treated with 1µM flg22. **E)** Comparison of the response to 1µM flg22 in Gerlach and *A. thaliana*. The data represent the means + SEM; n=8-16 discs in one biological experiment. The experiment was repeated three times independently with similar results. Asterisks (in Figure A) indicate that the mean value is significantly different from the control conditions (two-tailed Student’s t-test, ***p<0.001). Statistical differences between the samples (D) were assessed using a one-way ANOVA, with a Tukey honestly significant difference (HSD) multiple mean comparison post hoc test. Diferent letters indicate a signifcant diference, Tukey HSD, p<0.01. No letters in B, C mean that the data difference were not statistically significant.

### Flg22 and *Xcc*Flg22 interaction with Arabidopsis and putative *P. somniferum* FLS2

We identified a putative *PsFLS2* in the poppy genome based on the alignment of the protein sequences with AtFLS2 (Fig. S7), and we modelled its structure (Fig. S8).

Molecular docking analysis showed stronger binding affinity (Agu *et al*., 2023; Spassov, 2024) of *At*FLS2 to flg22 (Fig. 2A) compared to *Ps*FLS2 to flg22 (Fig. 2B). We used binding affinity to calculate *in silico* equilibrium dissociation constant (Kd) representing the ligand concentration required to occupy 50 % of the receptors (Hulme & Trevethick, 2010). *In silico* calculated Kd values were 0.0001 µM for *At*FL2-flg22 and 16.556 µM for *Ps*FLS2-flg22, showing that for *At*FLS2 the concentration to reach Kd is roughly 1.10^5^ lower than the concentration needed for reaching Kd in *Ps*FLS2 using flg22 as ligand (Table S2).

**Figure 2.**
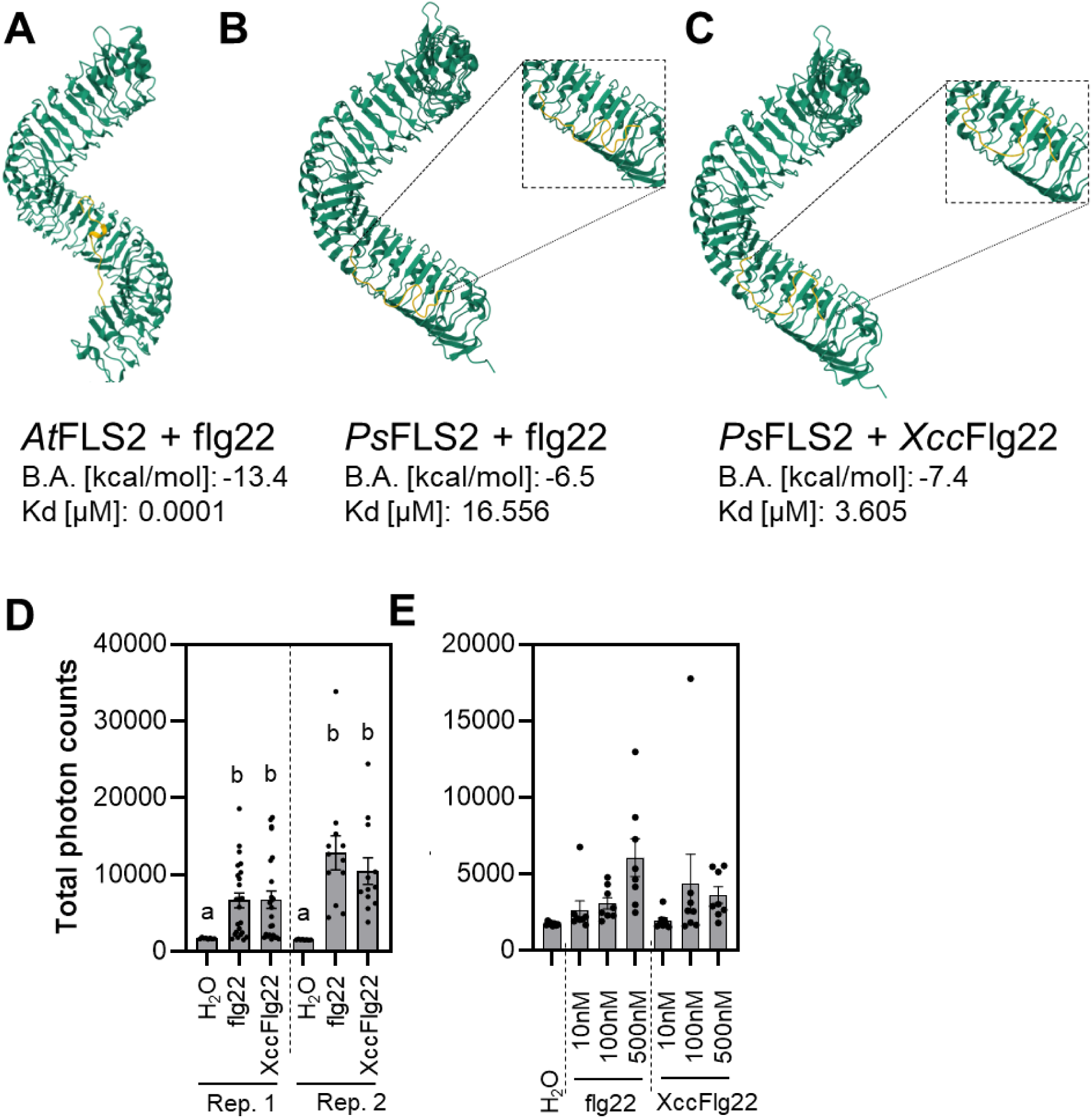
Response to flg22 originating from *Xanthomonas campestris* pv. *campestris*. **A)** Binding of *Pseudomonas aeruginosa* flg22 (flg22) to *Arabidopsis thaliana* FLS2 (*At*FLS2). **B)** Binding of *Pseudomonas aeruginosa* flg22 (flg22) to *Papaver somniferum* FLS2 (*Ps*FLS2). **C)** Binding of *Xanthomonas campestris* pv. *campestris* flg22 (*Xcc*Flg22) to *Papaver somniferum* FLS2 (*Ps*FLS2). B.A. – Binding affinity; Kd is estimated *in sillico*. **D-E)** Discs from Gerlach were cut from 5-6 week-old poppy plants. Discs were treated with 1 µM flg22 or *Xcc*Flg22 (D) and 10 – 500 nM flg22 or *Xcc*Flg22 (E). The data represent the means + SEM; n=8-16 discs in one biological experiment. The experiment was repeated two times independently with similar results. Statistical differences between the samples (D, E) were assessed using a one-way ANOVA, with a Tukey honestly significant difference (HSD) multiple mean comparison post hoc test. Different letters indicate a significant difference, Tukey HSD, P<0.01.

The flg22 (QRLSTGSRINSAKDDAAGLQIA), typically used in studies focused on plant immunity is derived from *Pseudomonas aeruginosa* (Trinh *et al*., 2023). However, *P. aeruginosa* is not a typical poppy bacterial pathogen, so we examined a variant, *Xcc*Flg22 (QRLSSGLRINSAKDDAAGLAIS), from poppy pathogen *Xanthomonas campestris* pv. *Campestris*. Molecular docking studies showed that *Xcc*Flg22 had a higher binding affinity for *Ps*FLS2 (Fig. 2C and Table S2), compared to flg22 and *Ps*FLS2 (Fig. 2B and Table S2). However, no significant difference between flg22 and *Xcc*Flg22 was observed in their ROS-inducing effects in poppy (Fig. 2D, E and S9). We decided to continue further PTI response analyses in poppy using flg22 peptide from *P. aeruginosa*.

### MAPK phosphorylation in *P. somniferum*

Activation of mitogen-activated protein kinase (MAPK) cascade, like the ROS burst, is a fast response triggered by the recognition of flg22 in plants (Frei dit Frey *et al*., 2014). We studied putative MAPK activation in poppy by using a pERK-antibody, which recognises phosphorylation of the canonical TEY motif in MAPKs, as a proxy (Yi *et al*., 2015) at two time points (15 and 30 minutes) following treatment (Fig. 3). Firstly, we started with infiltration using a needleless syringe using Gerlach as a model cultivar (Fig. 3A). As a result, we observed that wounding caused by infiltration of water also triggered putative MAPK activation with a similar intensity compared to infiltration with flg22 (Fig. 3A). To overcome this problem, we used the same approach as for ROS burst measurement using leaf discs incubated overnight in distilled water in the dark. Subsequently, we replaced the water with fresh water or with a solution containing flg22 without mechanically stressing the leaf discs. We observed an increased abundance of the protein recognised (putative MAPKs) by the pERK antibody upon flg22 treatment in the Gerlach cultivar (Fig. 3B). We used the leaf disc method to analyse all cultivars. In all of them, flg22 increased the abundance of pERK-recognised protein after 15 and 30 minutes (Fig. 3C, full blots in Fig. S10), likely corresponding to MAPKs phosphorylated at the TEY motif.

**Figure 3.**
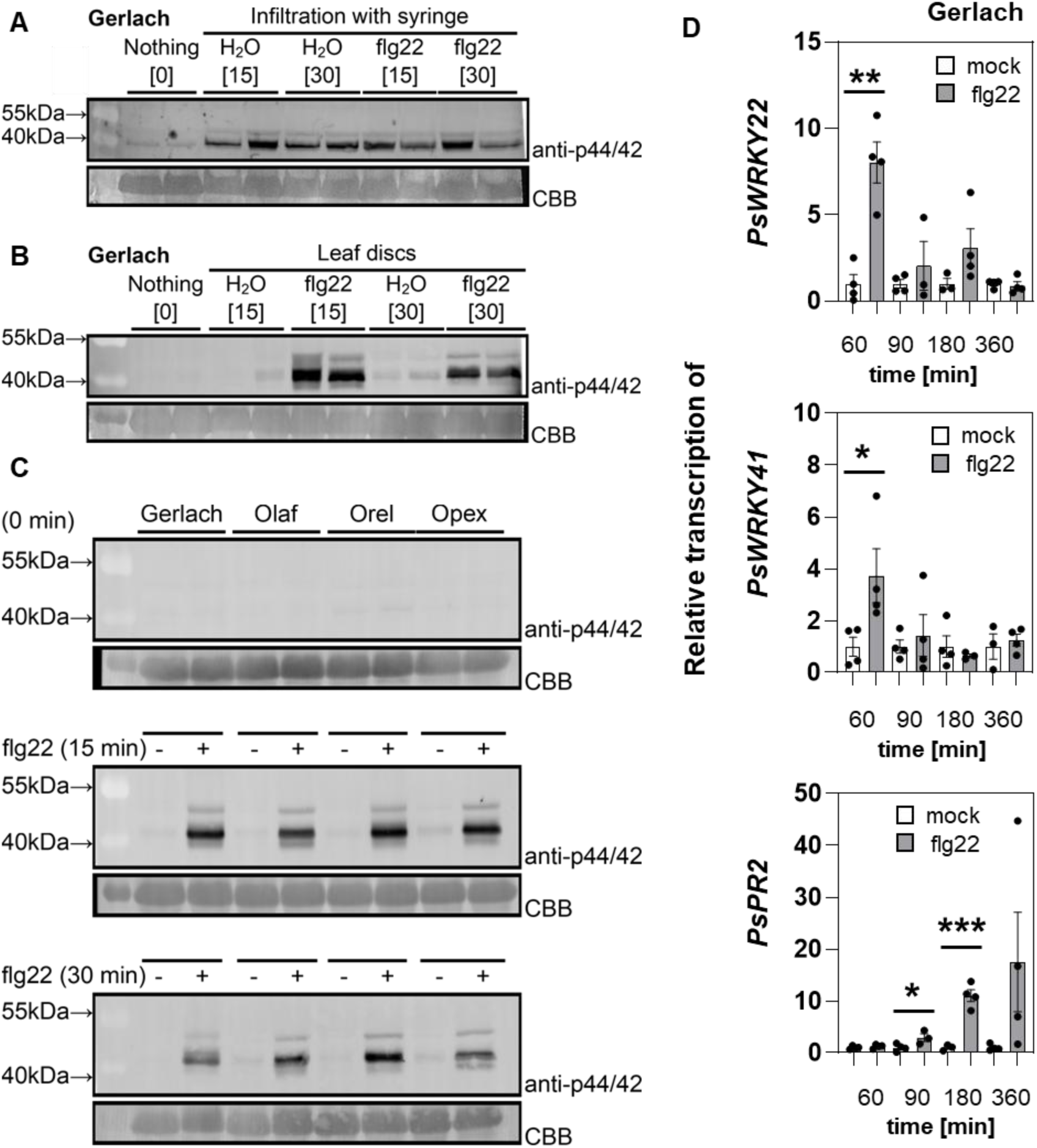
MAPK activation reflected by the abundance of pERK antibody recognised protein and gene transcription after treatment with flg22. The activation of the mitogen-activated protein kinases (MAPKs) was visualised by western blot analysis using the pERK phospho-p44/42 MAP kinase antibody. Comassie briliant blue (CBB) staining of the membrane was used as a loading control. **A)** Needleless syringae infiltration with 5µM flg22 (or water) of leaves from 5-6 week-old poppy plants. **B)** Treatment of leaf discs from 5-6 week old Gerlach plants with 5µM flg22 (or water). **C)** Treatment of leaf discs from 5-6 week old poppy plants (four genotypes) with 5µM flg22 (or water). **D)** The leaf discs were cut from 5-6 week-old poppy and treated with 5 µM flg22 for 60, 90, 180, 360 minutes. The relative transcription for controls (water treated samples) was normalised to mean 1. The data represent the means + SEM; n=3-4. Asterisks indicate that the mean value is significantly different from the control conditions (two-tailed Student’s t-test, *P<0.05, **P<0.01).

### Gene expression analysis of *P. somniferum*

We analysed gene expression changes after flg22 treatment to identify genes suitable for PTI monitoring in poppy. For that purpose, we used leaf discs to perform treatment similar to ROS burst and MAPK assay. We designed primers for orthologs of the genes connected with flg22-triggered responses in Arabidopsis (*PsWRKY22, PsWRKY33* and *PsFRK1*) (Zou *et al*., 2018; Bjornson *et al*., 2021), for flg22 receptor (*PsFLS2*) (Chinchilla *et al*., 2006) and for *PR2* gene (*PsPR2*), whose expression is sensitive to biotic and abiotic stress, but known to respond also to flg22 (Liu *et al*., 2023). Additionally, we studied the expression of three published putative poppy genes whose transcription was increased upon poppy treatment with bacterial endophyte *Microbacterium sp.* SMR1 (*PsCRK1*, *PsWRKY53*, *PsPRTS*) (Ray *et al*., 2021). We reannotated these three genes because the names were assigned based on a comparison with the annotated soybean genome. However, we used the available poppy genome (Guo *et al*., 2018), thus primers for *PsCRK1* target *PsCRK35, PsWRKY53* target *PsWRKY41* and *PsPRTS* target gene encoding the thaumatin-like protein. Among all the tested genes, *PsWRKY22*, *PsWRKY41* and *PsPR2* showed significantly enhanced transcript abundance following flg22 treatment in at least one analysed time point (Fig. 3D). *PsWRKY22* and *PsWRKY41* transcript levels were increased within 60 minutes and decreased over later time points (Fig. 3D), while *PsPR2* transcript abundance started to increase after 90 minutes and the increase continued through 180 minutes and 360 minutes (Fig. 3D). Other genes showed no significantly altered transcript levels after flg22 treatment (Fig. S11). However, we realised that using the discs for gene expression analyses is not ideal because we observed a significant effect on expression caused by cutting (Fig. S12). Although *PsWRKY22* exhibited clear induction of expression after flg22 treatment compared to mock samples, we also observed induction of *PsWRKY22* expression in mock samples compared to control (Fig. S12). *PsWRKY41* expression was strongly inhibited by cutting (Fig. S12). Our data suggest that *PsWRKY22*, *PsWRKY41* and *PsPR2* are promising candidate genes for monitoring PTI responses in poppy.

### Callose accumulation in *P. somniferum*

Compared to ROS burst and MAPK activation, which are transient and observed within minutes after PAMP or DAMP recognition, callose accumulation is generally detected within hours during PTI response (Kalachova *et al*., 2020; Mason *et al*., 2020; Liu *et al*., 2023). We used two approaches for flg22 treatment: (i) infiltration and (ii) leaf disc method. We analysed callose deposition 24 hours after treatment and observed similar callose accumulation levels in response to wounding (water infiltration in infiltrated samples, edge of the leaf discs) and flg22 (Fig. 4A-D). Callose accumulation following flg22 treatment in Arabidopsis is detectable and visible also in seedlings cultivated *in vitro* conditions (Gómez-Gómez *et al*., 1999; Zahid *et al*., 2017). This approach overcomes troubles with wounding. Thus, we used poppy seedlings cultivated in *in vitro* conditions and treated them with flg22. However, we did not observe any significant callose accumulation in poppy seedlings cultivated *in vitro* (Fig. 4E). These data indicate that wounding, not flg22, caused increased callose deposition.

**Figure 4.**
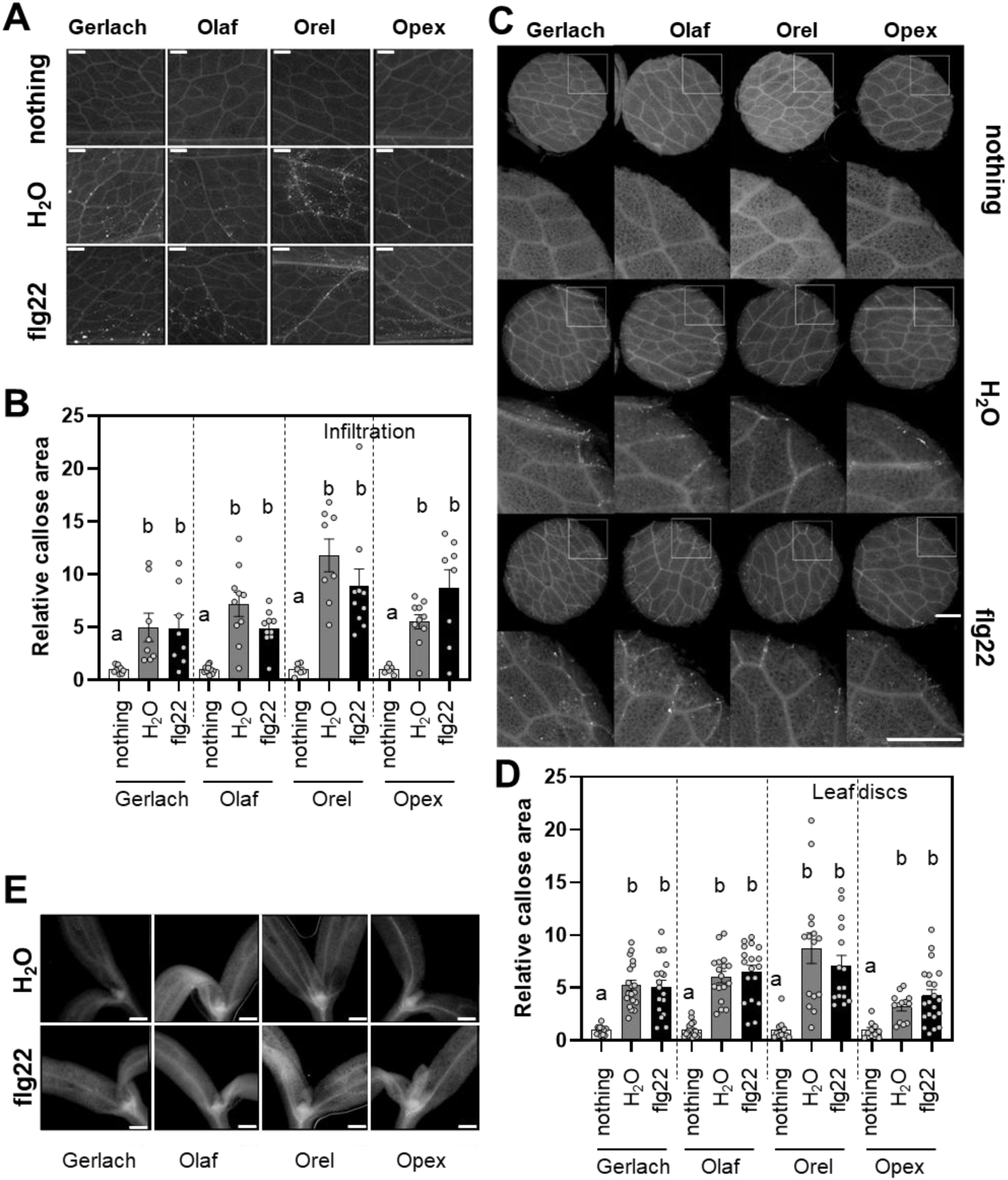
Callose accumulation. **A)** Representative images from 5-6 weeks old poppy leaves treated with 5µM flg22 (or H_2_O) using needleless syringae. **B)** Callose accumulation in 5-6 poppy leaves treated with 5µM flg22 (or H_2_O) using needleless syringae **C)** Representative images of discs and ROIs from 5-6 weeks old poppy leaves treated with 5µM flg22 (or H_2_O) using needleless syringae. **D)** Callose accumulation in discs cut from 5-6 week-old poppy plants. **E)** Representative images of 12-day-old poppy seedlings treated with 5µM flg22 (or water). The callose accumulation was monitored 24 hours after treatment. Bars represent 1 mm. The data represent the means + SEM; n=10-16. The experiment was repeated three times (B) or two times (C) independently with similar results. Statistical differences between the samples (B, C) were assessed using a one-way ANOVA, with a Tukey honestly significant difference (HSD) multiple mean comparison post hoc test. Different letters indicate a significant difference, Tukey HSD, P<0.01.

### Seedling growth inhibition and ROS burst in roots of *P. somniferum*

A typical long-term effect of plant immunity is the inhibition of plant growth, resulting from the so-called growth-defence trade-off (He *et al*., 2022). The screening is usually done with seedlings grown in liquid media *in vitro* (Gómez-Gómez *et al*., 1999; Janda *et al*., 2023). We used the same approach with poppy using liquid media containing sucrose (see Material and methods). Unlike Arabidopsis, flg22 did not significantly inhibit the growth of poppy seedlings (Fig. 5A and S13). Only the Olaf cultivar showed statistically significant growth inhibition, around 20 %, in four of six independent experiments (Fig. 5A and S13). However, we also tested the growth inhibition in liquid media without sucrose, and in such an experimental design, we observed significant growth inhibition in all cultivars in all independent experiments (Fig. 5A and S14). The inhibition was around 25 %, significantly weaker than Arabidopsis (Fig. 5A).

**Figure 5.**
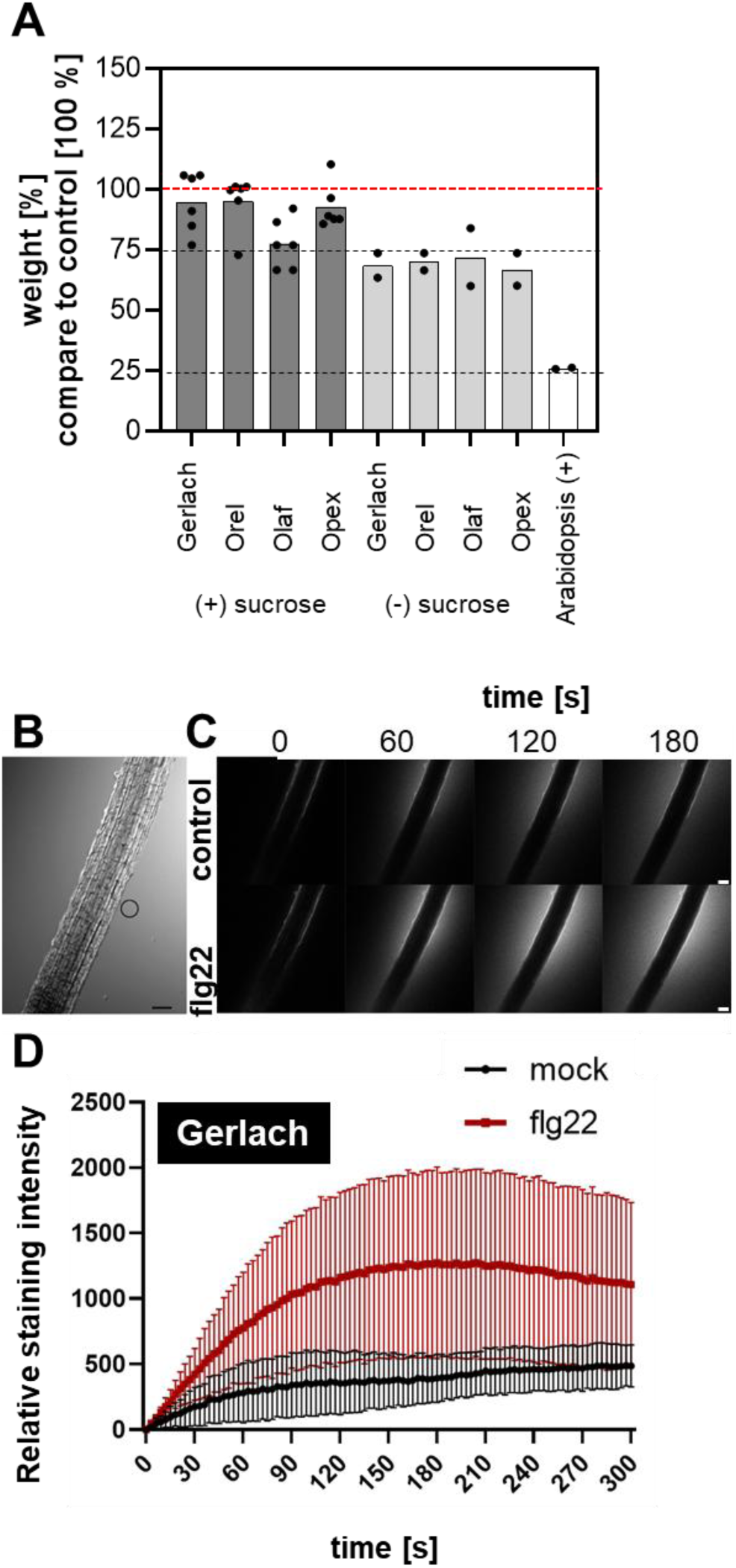
Seedlings growth inhibition and ROS production in roots. **A)** Growth inhibition analyses. [%] of 5µM flg22 treated seedlings weight compared to H_2_O (control) treated seedlings 5-7 days after the treatment in 1/2 MS medium +/- 1% (w/v) sucrose. The value of control was set as 100 % (represented by a red dashed line). The graph represents the mean of the means from independent biological experiments. Individual values represent the mean from one biological experiment in which was the weight of of 10-30 independent seedlings. The statistics for each individual independent biological experiment are provided in Figures S13 and S14. **B-D)** Five days-old seedlings of Gerlach cultivar were treated with 1µM flg22 B) Representative image (in the bright field) of the poppy root, depicting region is ROI where measurements were taken. **C)** Representative images of Amplex red signal (the halo around the root) caused by extracellular ROS burst after 1µM flg22 treatment. Scalebar = 100 µm. **D)** Analysis of Amplex red and ROS accumulation caused fluorescence in root elongation zone upon 1µm flg22 n≥4, mean with SD.

Observing no callose (Fig. 4E; ROS Fig. S15) or weaker (growth inhibition) responses in *in vitro* experimental conditions, we searched for another possible screening method for PTI analysis in seedlings. For that purpose, we used the method developed by Kulich et al. (2025) to analyse apoplastic ROS burst in roots using Amplex Red dye. The reaction provides red fluorescence upon oxidation to resorufin in the presence of H_2_O_2_. The advantage of the method is that it monitors very fast response and enables to have as control and treated sample the same seedling. We observed clear production of ROS (halo around the poppy roots Fig. 5B, C) in apoplast within 5 minutes after treatment with flg22 (Fig. 5D).

### Ethylene and salicylic acid production in *P. somniferum*

Production of ethylene, a gaseous defence-related phytohormone, is a fast and rapid response to flg22 recognition in Arabidopsis (Felix *et al*., 1999). Using a needleless syringe, we treated poppy leaves with 5µM flg22 and distilled water as mock control. We repeatedly observed significantly increased ethylene production as a response to wounding, which was caused by the infiltration of the H_2_O in the Gerlach cultivar (Fig. 6A). In Opex and Orel, the trend was similar, but the difference was not significant. In the Olaf cultivar, we observed a significant difference between H_2_O and flg22 treatment (Fig. 6A).

**Figure 6.**
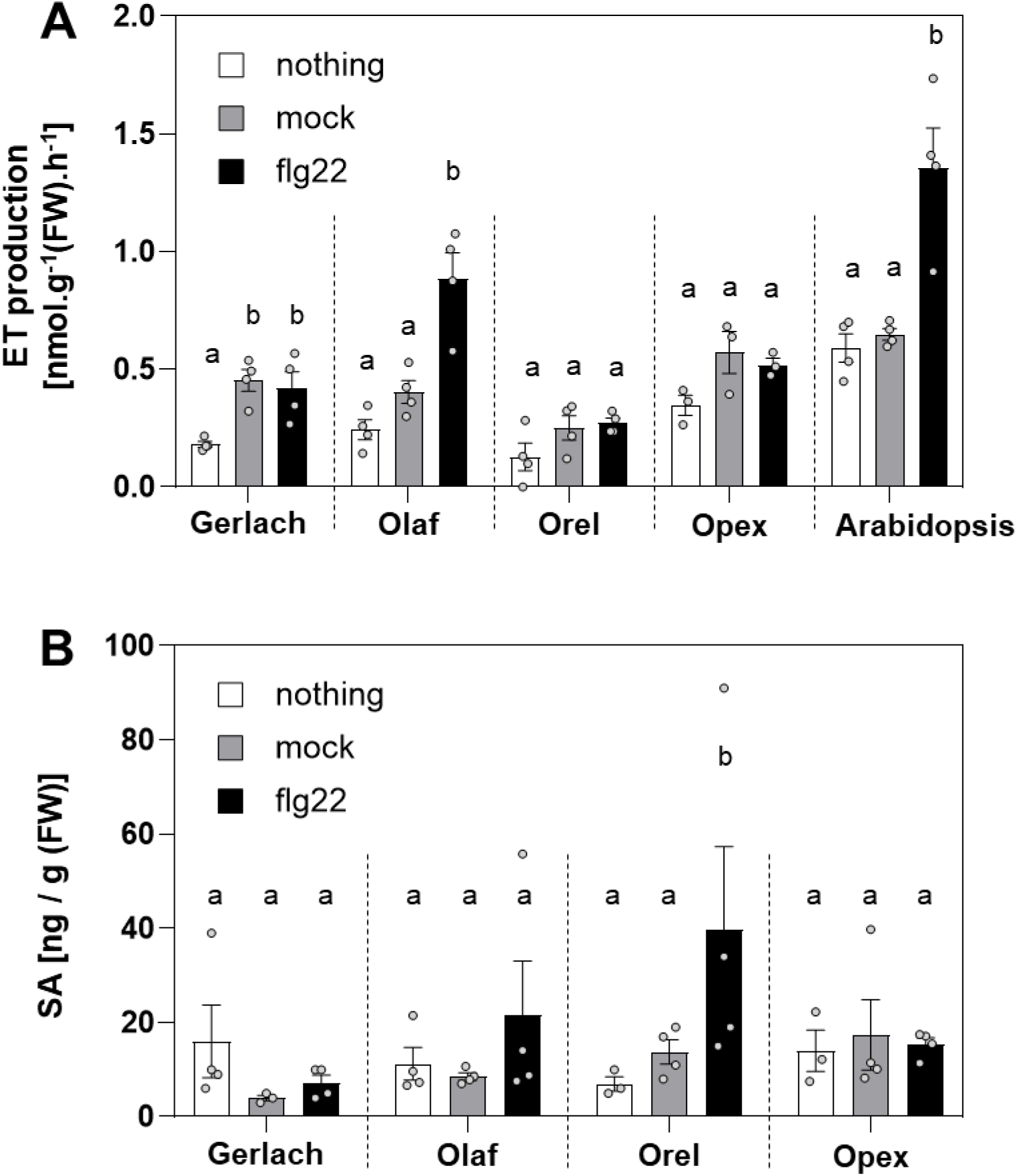
Ethylene and salicylic acid accumulation. The leaves from 5-6 week-old poppy and *A. thaliana* plants were treated with 5 µM flg22 with infiltration using a needleless syringe. As a control, non-treated leaves and leaves treated with water were used. **A)** The ethylene production from poppy and Arabidopsis leaves was analysed 4 hours after treatment. **B)** Salicylic acid (SA) concentration was analysed 24 h after the treatment with flg22. The data represent the means + SEM; n=3-4 (ethylene) and n=4 (salicylic acid). Statistical differences between the samples among one genotype were assessed using a one-way ANOVA, with a Tukey honestly significant difference (HSD) multiple mean comparisons post hoc test. Diferent letters indicate a significant difference, Tukey HSD, p<0.01.

Additionally, we analysed another typical defence-related phytohormone, salicylic acid (SA). For that purpose, we used UPLC with a fluorescence detector inspired by a method published in a study measuring SA in poppy after endophyte inoculation (Ray *et al*., 2021). However, we were unable to detect any SA. Thus, we used the method we previously successfully used for Arabidopsis analyses (Pluhařová *et al*., 2019). With this method, we identified SA in the range from 5-40 ng / g of fresh weight tissue (Fig.6B). However, we did not observe any effect caused by wounding. Only for Orel cultivar caused the treatment with flg22 showed a significant increase in SA concentration (Fig. 6B). The SA content in poppy was low compared to our Arabidopsis data (around 1-2 µg / g FW) (Hao *et al*., 2012; Pluhařová *et al*., 2019) or levels observed in other plant species (0,3-1 µg/g FW in *Humulus lupulus*, 1-3 mg / g FW in willow bark) (Petrek *et al*., 2007; Patzak *et al*., 2013).

## DISCUSSION

Research on *Papaver somniferum* L. (poppy) has been so far predominantly focused on biosynthetic pathways and the production of secondary metabolites (Singh *et al*., 2019). It was shown that poppy, in particular *Papaver rhoeas*, is suitable as a model for studying cell death, especially in pollen (Geitmann *et al*., 2004; Wilkins *et al*., 2015). By contrast, only a few studies were focused on the molecular aspects of the interactions between poppy and its pathogens (Thangavel *et al*., 2020; Ray *et al*., 2021) although poppy yield is strongly influenced by pathogens and pests (Bailey *et al*., 2000; Thangavel *et al*., 2018). To the best of our knowledge, no study has been available on plant immunity in poppy. Here, we provide the first comprehensive investigation of PTI in poppies.

As a base cultivar for analysing breadseed poppy PTI responses, we selected the spring blue seed cultivar Gerlach, which was introduced into the market in 1990. Using Gerlach, we showed that among all tested peptide elicitors, flg22 is the most potent peptide to induce ROS burst (Fig. 1). However, flg22-triggered ROS production was significantly lower in Gerlach compared to Arabidopsis (Fig. 1E). This could be explained by a lower binding affinity of *Ps*FLS2 to flg22 compared to the binding affinity of *At*FLS2 (Fig. 2A). We showed that in poppy flg22 induced putative MAPK phosphorylation (Fig. 3B, C), seedling growth inhibition (Fig. 5A), ROS burst in roots (Fig. 5B) and we identified candidate genes for monitoring PTI (Fig. 3D). Surprisingly, compared to other plant species (Tsuda *et al*., 2008; Nguyen *et al*., 2010; Skottke *et al*., 2011; Lloyd *et al*., 2014), no callose accumulation (Fig. 4), ethylene (Fig. 6A) and SA (Fig. 6B) production was observed after flg22 in Gerlach.

We compared the PTI responses of Gerlach with other poppy cultivars which were selected to represent the common breadseed poppies on the fields in Central Europe: spring white seed (Orel), overwintering blue seed (Olaf) and another spring blue seed (Opex). Most of the PTI responses were similar in all cultivars compared to Gerlach, with few exceptions. In contrast to Gerlach, we observed increased SA concentration in Orel in response to flg22 (Fig. 6B). Interestingly, SA concentration in poppy cultivars is at least ten times lower than in other plant species (Hao *et al*., 2012). Thus, deciphering the role and potential of SA in poppy defence against pathogens should be further investigated. For example, SA signalling can be monitored with transcriptomic analysis or testing of the SA treatment on poppy growth and defence against pathogens. Unlike other cultivars, Olaf was the only cultivar in which flg22 triggered a significant increase in ethylene production (Fig. 6A). Olaf also exhibits growth inhibition after flg22 treatment in media containing sucrose (Fig. 5A) and has a slower induction of ROS production (Fig. S5). Spring cultivars might have different strategies for defence compared to overwintering cultivars. Having more spring and overwintering blue seed cultivars in germline bank opens research opportunities which might be unique among other plant species.

Analysis of MAPK phosphorylation and callose accumulation revealed that wounding stress is a challenge in PTI investigation in poppy, e.g. in comparison to studies performed on *Brassicaceae* or *Solanaceae*, in which syringe infiltration was successfully used (Nguyen *et al*., 2010; Lloyd *et al*., 2014). In poppies using syringe infiltration, we observed the same level of putative MAPK phosphorylation for water-treated samples as for flg22-treated ones (Fig. 3A). Using a leaf discs approach, we overcame this problem (Fig. 3B). Monitoring callose, we observed the significant callose accumulation after syringe infiltration of water at the same level as for flg22-treated samples in the site of infiltration (Fig. 4). Additionally, using leaf discs approach we observed callose accumulation near to the cut region, but we did not see any difference between water and flg22-treatment (Fig. 4). The question is if not observed difference between water and flg22-treatment is because poppy may not include callose accumulation among PTI responses or because we did not find optimal conditions to study callose accumulation after flg22 treatment in poppy.

To overcome the problem with wounding, we used seedlings grown *in vitro* for analysis (Fig. S2), but we did not see callose accumulation in seedlings (Fig. 4) even though we did not observe ROS burst in seedlings using the luminol-based method (Fig. S15). Nevertheless, we used a recently developed assay for monitoring extracellular ROS burst in seedlings roots using Amplex Red fluorescent dye (Kulich *et al*., 2025). With this approach, we observed clear induction of ROS burst caused by flg22 treatment in roots (Fig. 5B, C). This method seems to have great potential for screening the root reaction to molecular patterns, and it would be great to incorporate it among the set of methods for PTI studies, especially in poppy. Additionally, in seedlings, we observed growth inhibition after flg22 treatment. To note the reproducible inhibitory effect of flg22, we observed just using a medium without sucrose (Fig. 5A). This is the methodologically relevant observation that such a compound as sucrose, which is commonly used in the Arabidopsis research (Wierzba & Tax, 2016; Janda *et al*., 2023), can abolish the effects of flg22 in poppy.

The response of poppy to wounding represents the opportunity for deeper analysis. PTI is interconnected with wounding, sharing similar responses on the molecular level (Choi & Klessig, 2016). PAMPs represent bacterial molecules inducing immunity, but during infection, plants produce DAMPs, molecules whose production is triggered by damage of plant tissue (Hou *et al*., 2019). Our study used one DAMP, *At*Pep1, peptide originating from Arabidopsis (Huffaker *et al*., 2006). Based on the available literature, *At*Pep1 might be specific for Arabidopsis (*Brassicaceae*). However, we monitored the weak induction of ROS burst after *At*Pep1 treatment in poppy leaves (Fig. 1, S3). Investigating the presence of DAMP peptides similar to pep1 in poppy will be interesting.

We showed that with established methods, it would be possible to perform systematic screenings of poppy PTI responses to other known elicitors, such as chitin, to which the immune response is even more conserved in plants than to flg22 (Gimenez-Ibanez *et al*., 2019) or for searching for novel elicitors of poppy immunity. The obtained knowledge would streamline future poppy breeding for better resistance to pathogens. For example, the information that poppy does not respond to known PAMPs or DAMPs with a known receptor in other plant species could potentially enable us to design a way to obtain novel transgenic poppy resistant to a particular pathogen using an approach similar to the introduction of EFR receptor into plant species from distinct families than *Brassicaceae* (Lacombe *et al*., 2010; Schwessinger *et al*., 2015; Schoonbeek *et al*., 2015; Lu *et al*., 2015; Boschi *et al*., 2017; Mitre *et al*., 2021; Piazza *et al*., 2021; Adero *et al*., 2023). However, for the success of this approach, existing strict legal regulations and negative public opinions create significant obstacles (Garcia-Alonso *et al*., 2022). Additionally, screening of the PTI responses in available poppy cultivars e.g. from Czech poppy seeds bank (https://grinczech.vurv.cz) (REF) or performing EMS mutagenesis analysis would further enable deciphere PTI signalling in poppy.

## CONCLUSION

Our study provides a methodological pipeline for studying PTI and illustrates its complexity in poppy. Activated PTI after flg22 treatment is evident, as suggested by ROS burst, putative MAPK phosphorylation, seedling growth inhibition, and altered gene transcription. However, ethylene and SA production also increased in certain cultivars. Interestingly, callose accumulation seems to be independent of flg22 treatment. Our results indicate that attention must be paid to overlaps between PTI and wounding when investigating PTI. The insights and methodological know-how gained from this research not only advance our understanding of plant immunity in poppy but also pave the way for future studies to improve disease resistance in poppy. Further research should explore the genetic basis of the observed variability and investigate the role of additional signalling molecules to develop comprehensive strategies for enhancing the resilience of poppy against various pathogens.

## ACKNOWLEDGEMENTS

We thank Petra Fialová for her excellent support and Prof Vladislav Čurn (USB) for fruitful discussions. We thank Natálie Hradecká, who helped with the first poppy experiments in the MPMI Lab. We thank Dr Hana Leontovyčová for helping with the preparation of the samples for salicylic acid measurement. We thank the Institute of Molecular Plant Biology, Biology Centre, particularly Jan Kadlec, for excellent support. Part of the work was carried out with the support of the Growth Facility (BC Core Facilities; IPMB BC CAS; supported by Horizon Europe, MOLIPEC, ID 101087030). We acknowledge the core facility LMH, the BC CAS supported by the MEYS CR (LM 2023050 Czech-BioImaging). We want to acknowledge Grammarly, which we used for the improvement of clarity, flow, and grammatical accuracy.

This research was supported by MEYS (from the EU Operational Programme), the nr. CZ.02.2.69/0.0/0.0/18_053/0016975 (MJ), by MEYS Inter -Excellence II programme (project LUC23146, MSMT – 1942/2023-2) (JSHO, OI, OH, MJ), by exRNA-PATH COST action CA20110 (JSHO, MJ), by Team GaJU support from Faculty of Science, University of South Bohemia in České Budějovice (JSHO), support from GACR 23-04866S (MW, AZ), support from European Union, Horizon Europe MOLIPEC, ID 101087030 (IK), by Research Infrastructure METROFOOD-CZ supported by the Ministry of Education, Youth, and Sports of the Czech Republic, project No. LM2018100 (PM).

## AUTHOR CONTRIBUTIONS

JSHO – ROS analyses, bioinformatics-*Ps*FLS2 and *Xcc*flg22, growth inhibition, sample preparation for immunoblot, SA, gene expression, callose, writing of the manuscript, OI – callose analysis, growth inhibition, ROS measurement in poppy roots samples preparation for MAPK, gene expression and SA analyses; AZ – design and immunoblot analysis, writing of the manuscript; MH – gene expression analysis, JK – ethylene measurements; BK – ROS analyses, growth inhibition; IV - contributed to establishment of growing of the poppy and to callose experiments; SP – performed seedlings growth inhibition experiment using MS medium without sucrose; PM – SA analysis; MM – contributed to establishment of growing of the poppy, *P. somniferum*, primers design; PF – SA analysis; AR – poppy seeds multiplication and storage, establishment of poppy cultivation, OH – establishment of poppy cultivation, growth inhibition analyses; IK – ROS measurement in poppy roots; MW – design of immunoblot experiment, writing of the manuscript; MJ – conceptualization of the study, ROS analyses, preparation of samples for ethylene analyses, writing and finalising of the manuscript. All the authors commented on the manuscript before its finalising and approved the final version.

