## Supplementary material for "Pattern-triggered immunity in blue and white seed cultivars of *Papaver somniferum*"

*Hernández Orozco et al. 2025*

**Figure S1.** Representative photos of *Papaver somniferum* plants growth in soil from 5$^{th}$ to 30$^{th}$ day.

**Figure S2.** Representative figures of *Papaver somniferum* growth in sterile conditions from seed germination to 10$^{th}$ day.

**Figure S3.** ROS burst in poppy after treatment with elicitors.

**Figure S4** Positive control for triggering ROS burst by the peptides in distinct potatoes and Arabidopsis.

**Figure S5.** ROS burst in poppy cultivars triggered by treatment with flg22.

**Figure S6.** ROS burst in Gerlach and Arabidopsis triggered by treatment with distinct concentrations of flg22.

**Figure S7**. Alignment of *At*FLS2 (AT5G46330.1) and *Ps*FLS2(XP_026454075.1) protein sequences.

**Figure S8.** Confidence plots of 3D *Ps*FLS2 predicted structure.

**Figure S9.** ROS burst in Gerlach triggered by treatment with distinct flg22.

**Figure S10.** Original documentation of full gels of immunoblots which were shown in figure 3.

**Figure S11** Gene expression after treatment with flg22.

**Figure S12**. Gene expression after treatment with $H_2O$ compared to non-treated samples.

**Figure S13** Raw data for seedlings growth inhibition in media containing sucrose shown in figure 5.

**Figure S14** Raw data for seedlings growth inhibition in media without sucrose shown in figure 5.

**Figure S15** ROS burst in seedlings of four poppy cultivars after flg22 treatment.

**Table S1.** Primers used for RT-qPCR in *P. somniferum*

**Table 2.** List of binding affinity energies for *P. somniferum* and *A. thaliana* FLS2 and flg22 and *Xcc*Flg22

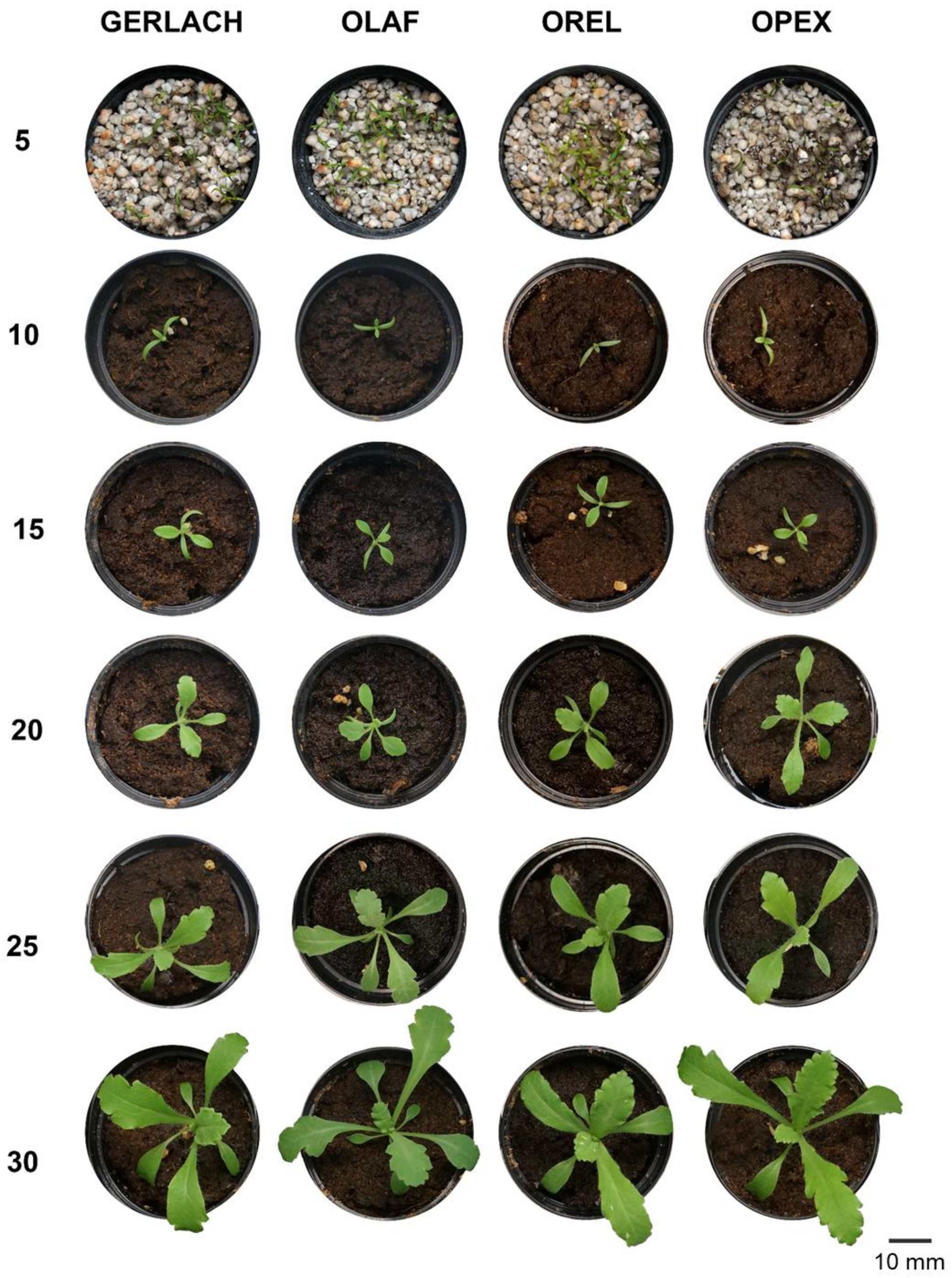


**Figure S1.** Representative photos of *Papaver somniferum* plants growth in soil from 5$^{th}$ to 30$^{th}$ day.

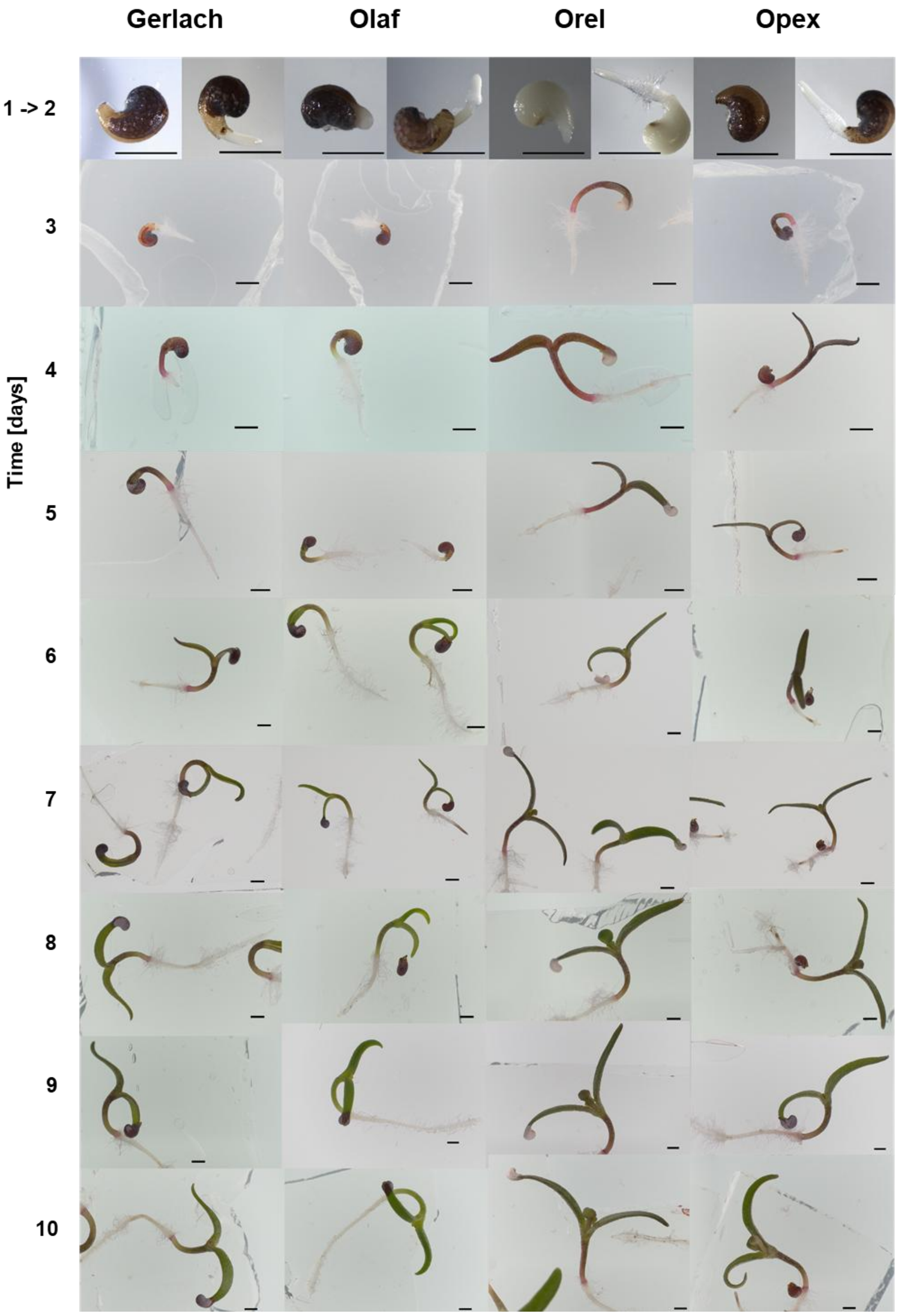


**Figure S2.** Representative figures  of *Papaver somniferum* growth in sterile conditions from seed germination to 10$^{th}$ day.

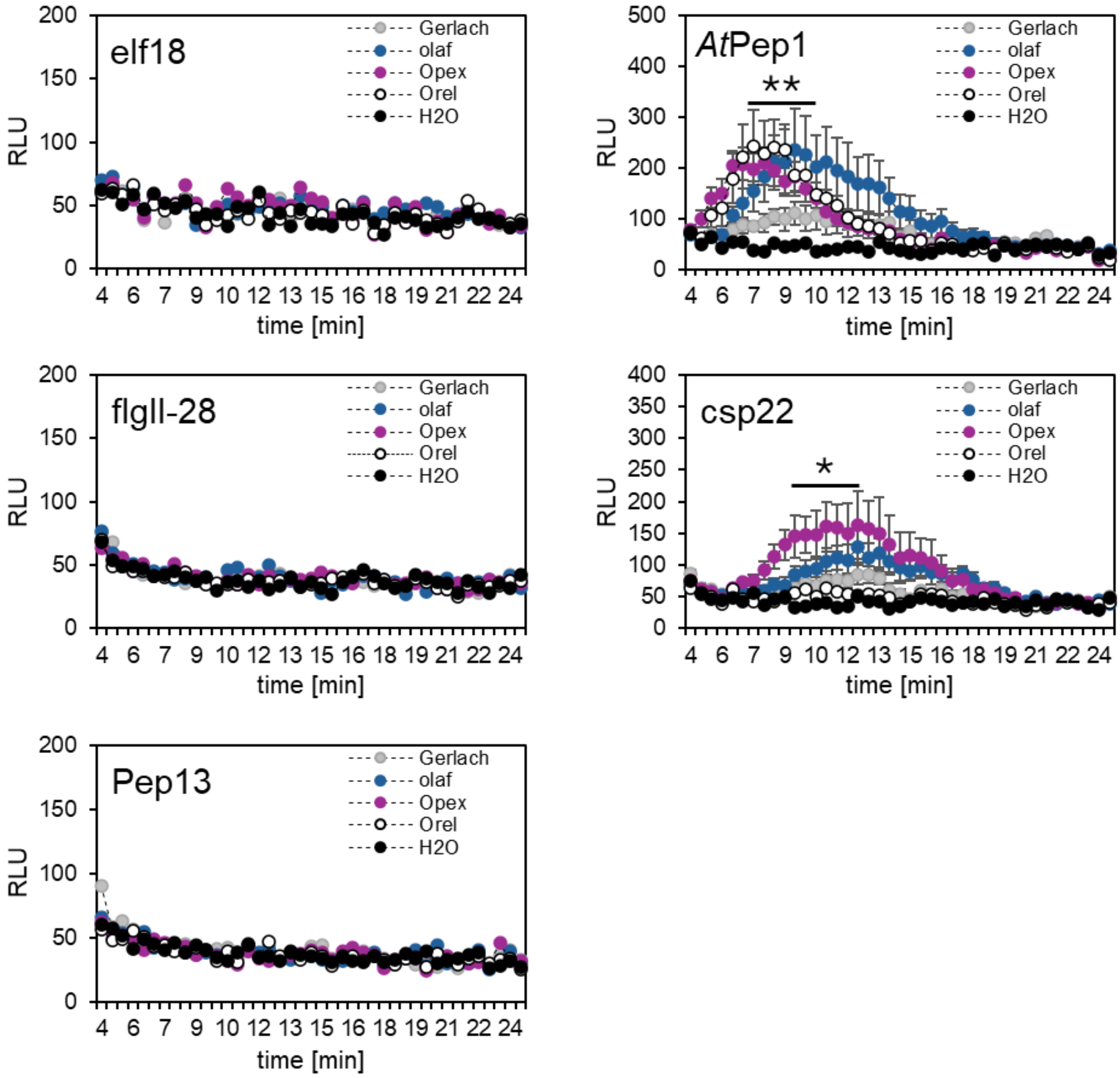


**Figure S3. ROS burst in poppy after treatment with elicitors.** Discs were cut from 5-6 week old poppy plants. Discs were treated with 1 µM: flgII-28, csp22, *At*Pep1, elf18, Pep13. The data represent the means; n=8-16 discs in one biological experiment. The experiment was repeated three times independently with similar results. Asterisks indicate that the mean value is siginificantly different from the control ($H_2O$) conditions (two-tailed Student´s t-test $^*p<0.05$, $^{**}p<0.01$).

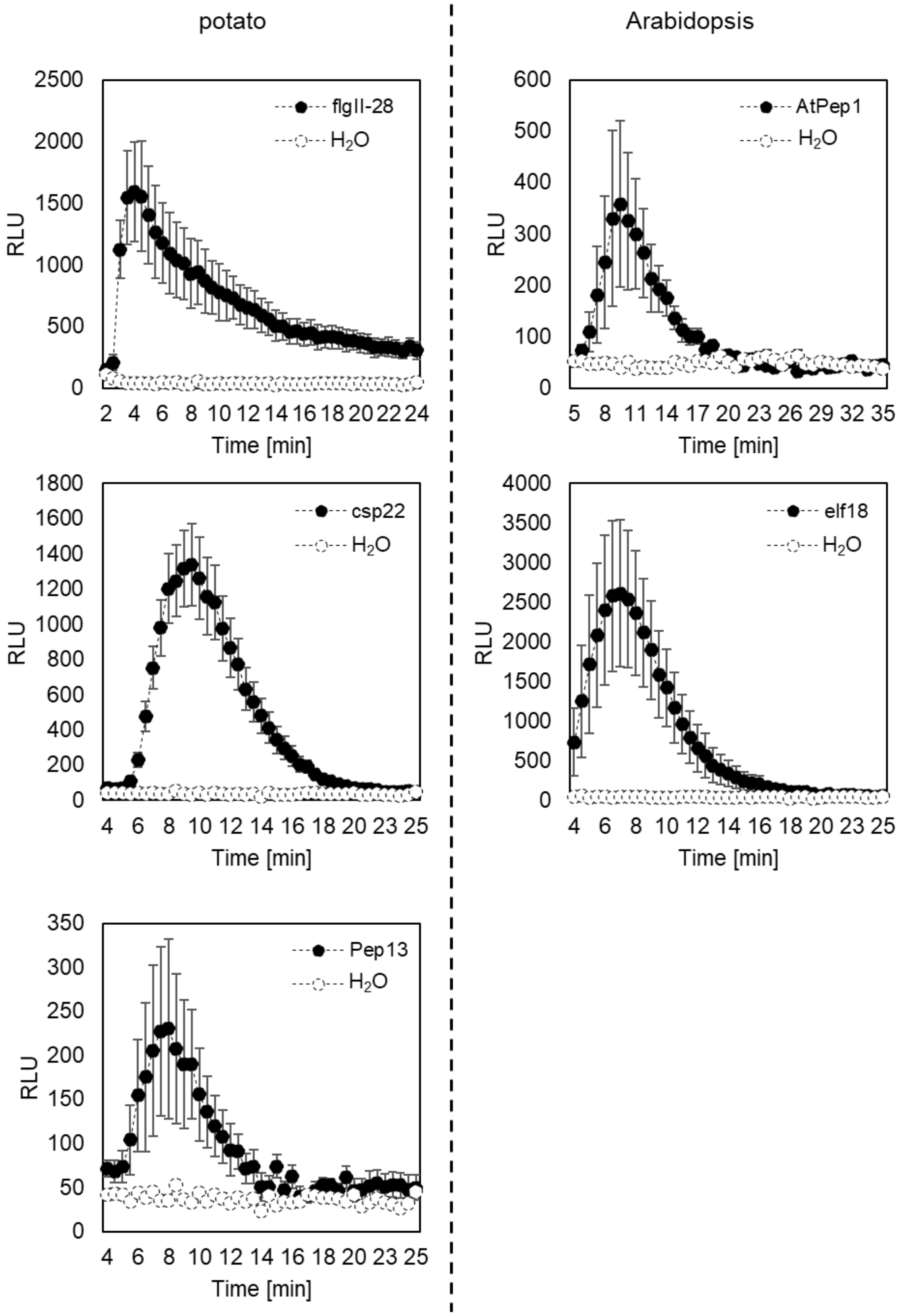


**Figure S4 Positive control for triggering ROS burst by the peptides in distinct potatoes and Arabidopsis.** Discs were cut from potato (Pep13, csp22, FlgII-28), Arabidopsis (elf18, *At*Pep1). The data represent mean and SEM from 8-12 discs. The experiment was repeated two times with similar results.

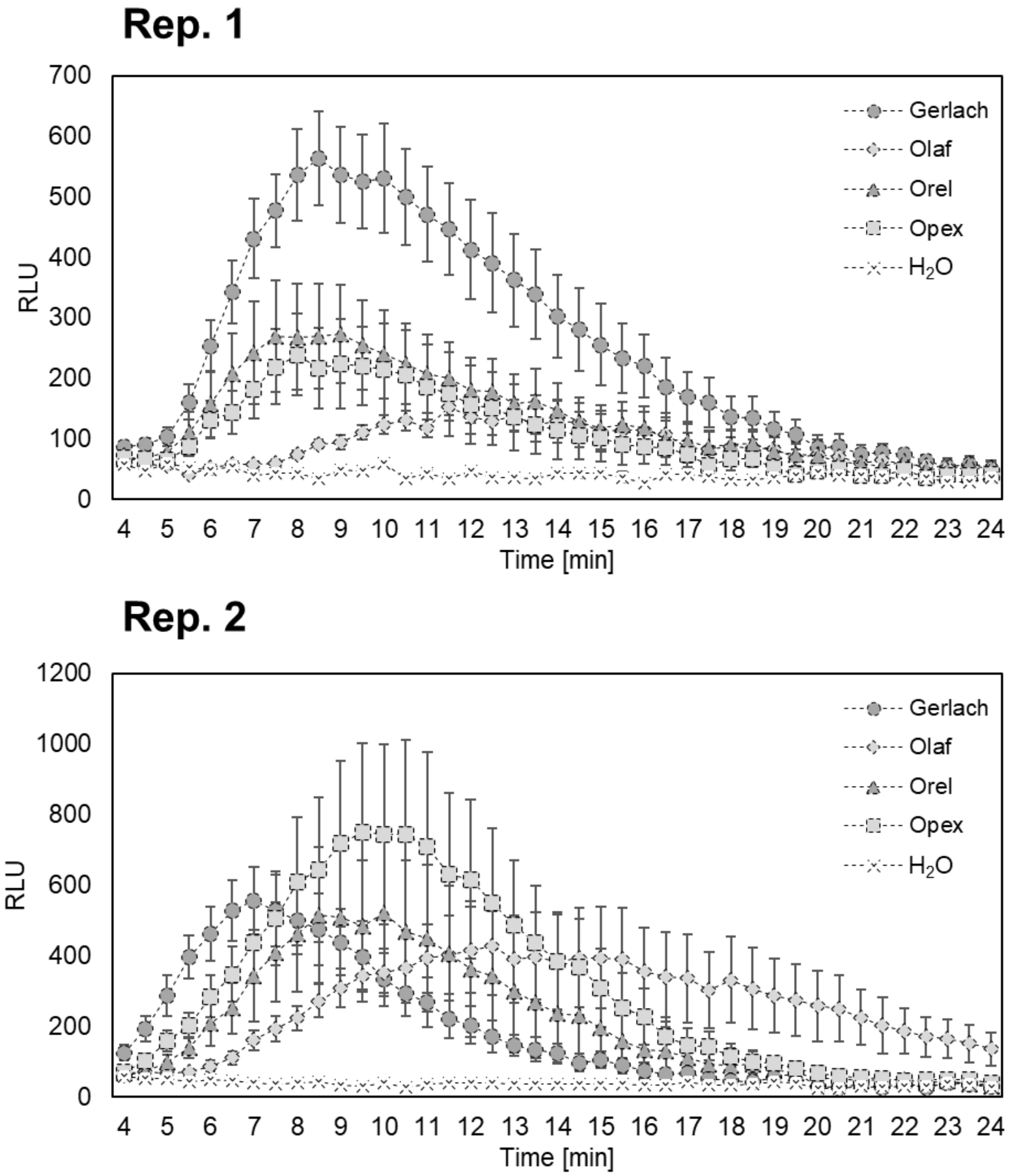


**Figure S5. ROS burst in poppy cultivars triggered by treatment with flg22.** Discs were cut from 5-6 week old poppy. The data represent the means and SEM; n=8-16 discs. Two biological experiments are shown.

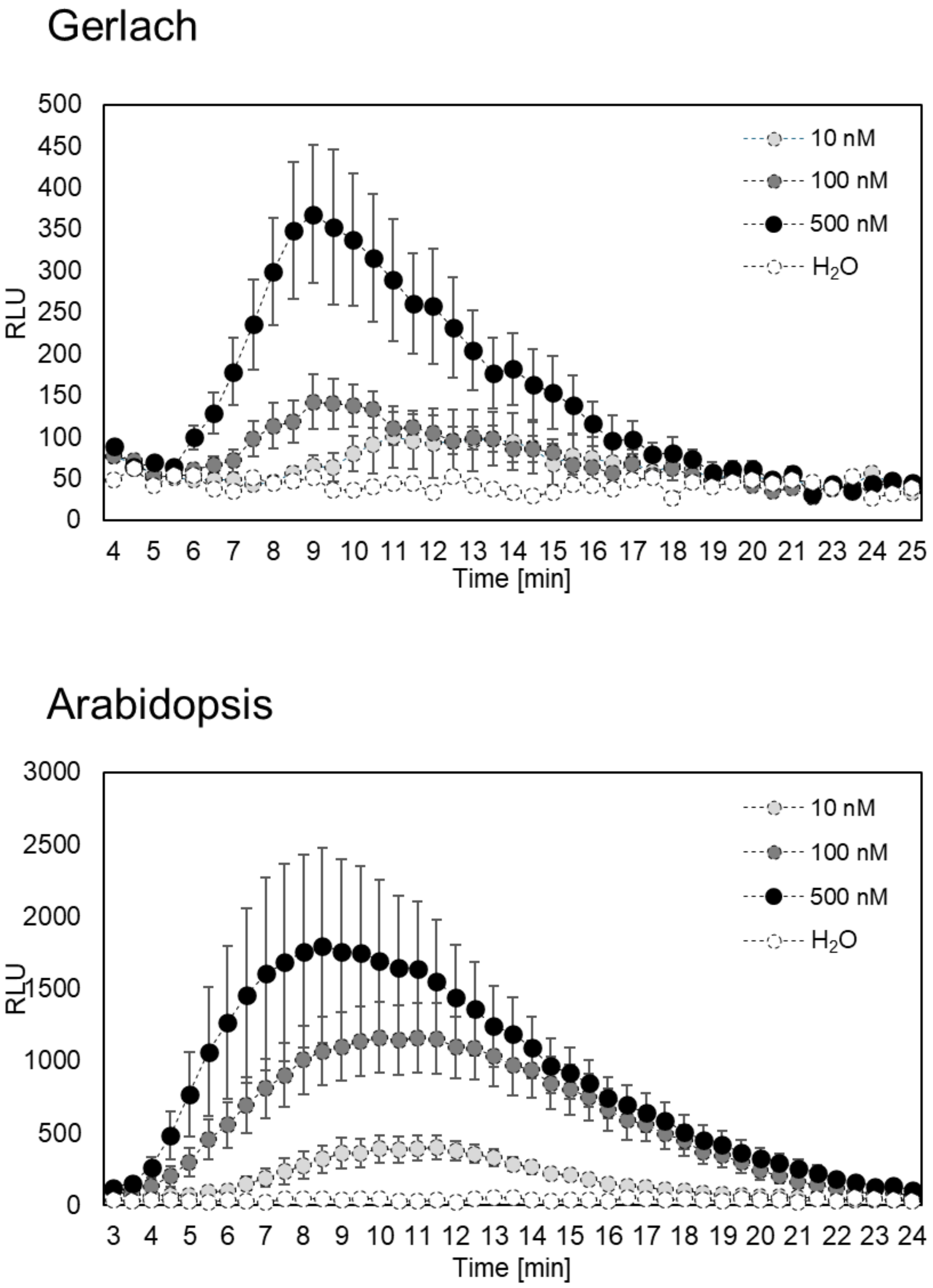


**Figure S6. ROS burst in Gerlach and Arabidopsis triggered by treatment with distinct concentrations of flg22.** Discs were cut from 5-6 week old Gerlach and *A. thaliana*. The data represent the means and SEM; n=8-16 discs. The experiment was repeated in two biological experiments with similar results.

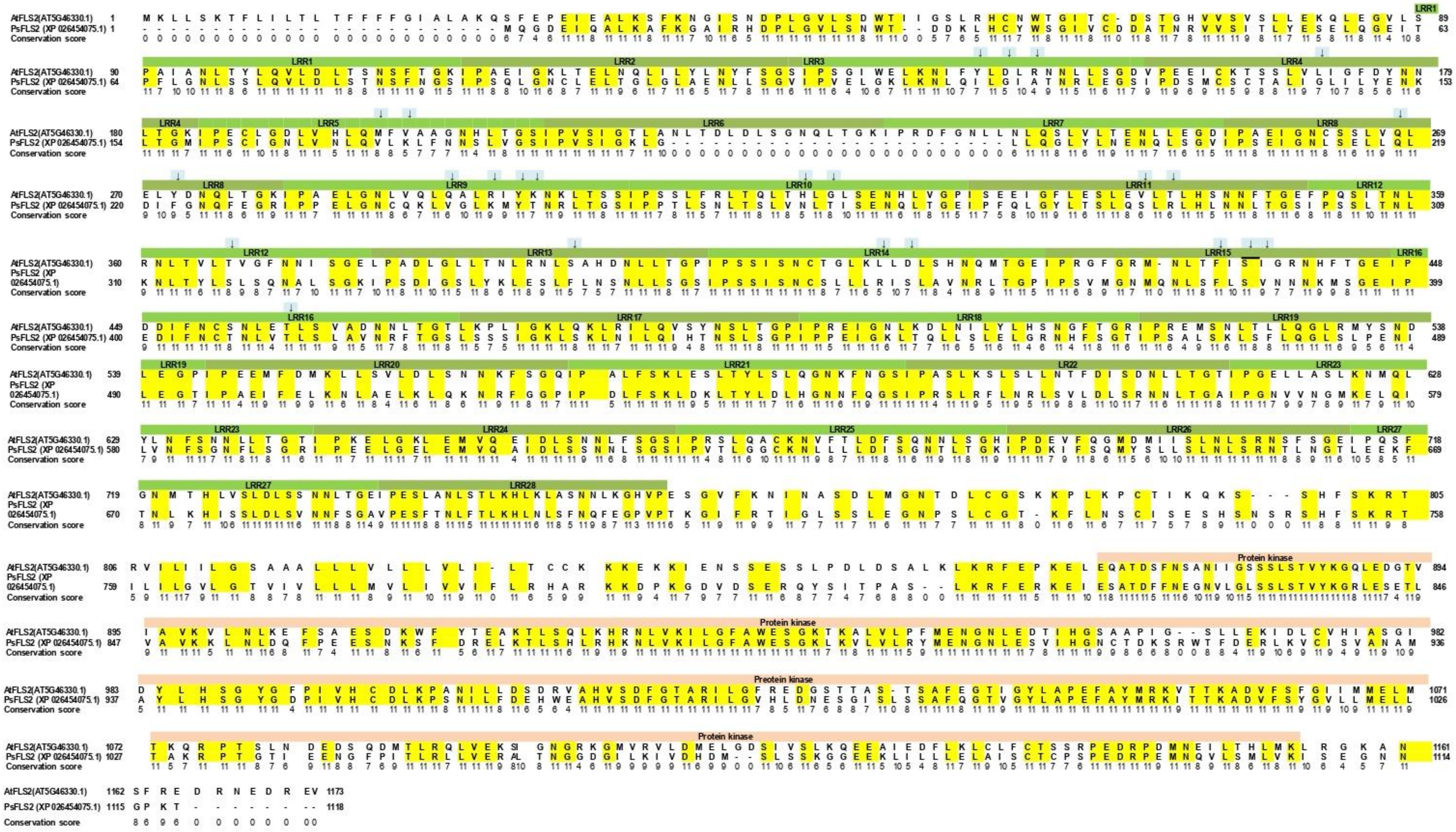


**Figure S7. Alignment of *At*FLS2 (AT5G46330.1) and *Ps*FLS2(XP_026454075.1) protein sequences.** Yellow color represent the conserved amino acid regions, LRRs are represented with green, light orange represents the protein kinase domain, and potential binding amino acids with flg22 are pointed with an arrow on light blue background.

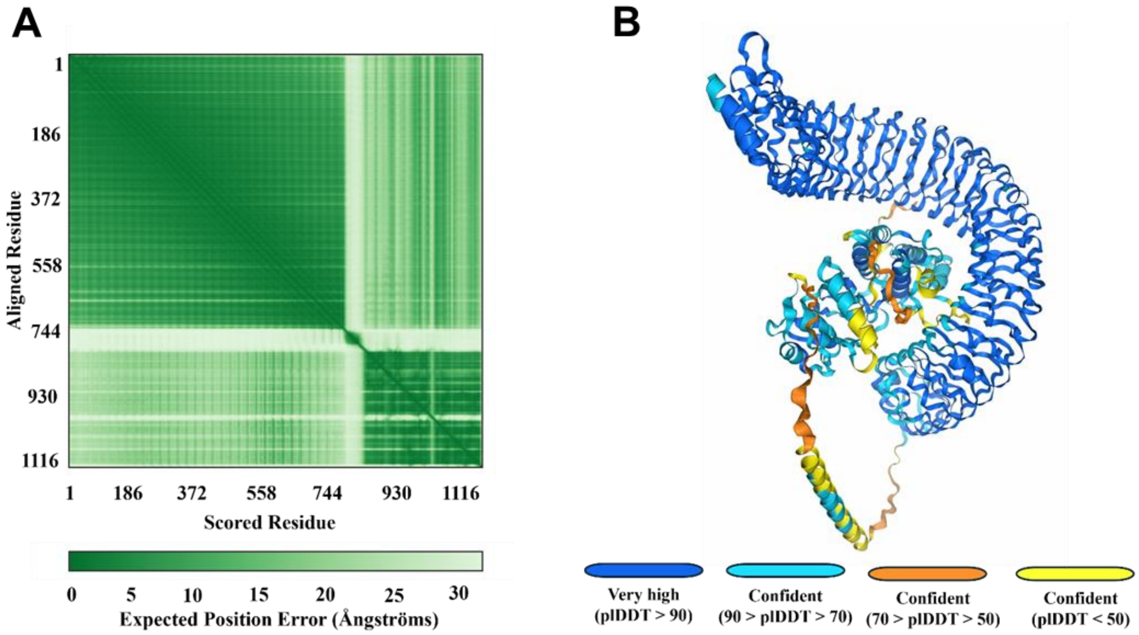


**Figure S8. Confidence plots of 3D *Ps*FLS2 predicted structure. A)** Predicted Aligned Error (PAE) plot for predicted *Ps*FLS2. **B)** Predicted local Distance Difference Test **(**pLDDT). Both plots were generated automatically on AlphaFold2 server version 2.3.0.

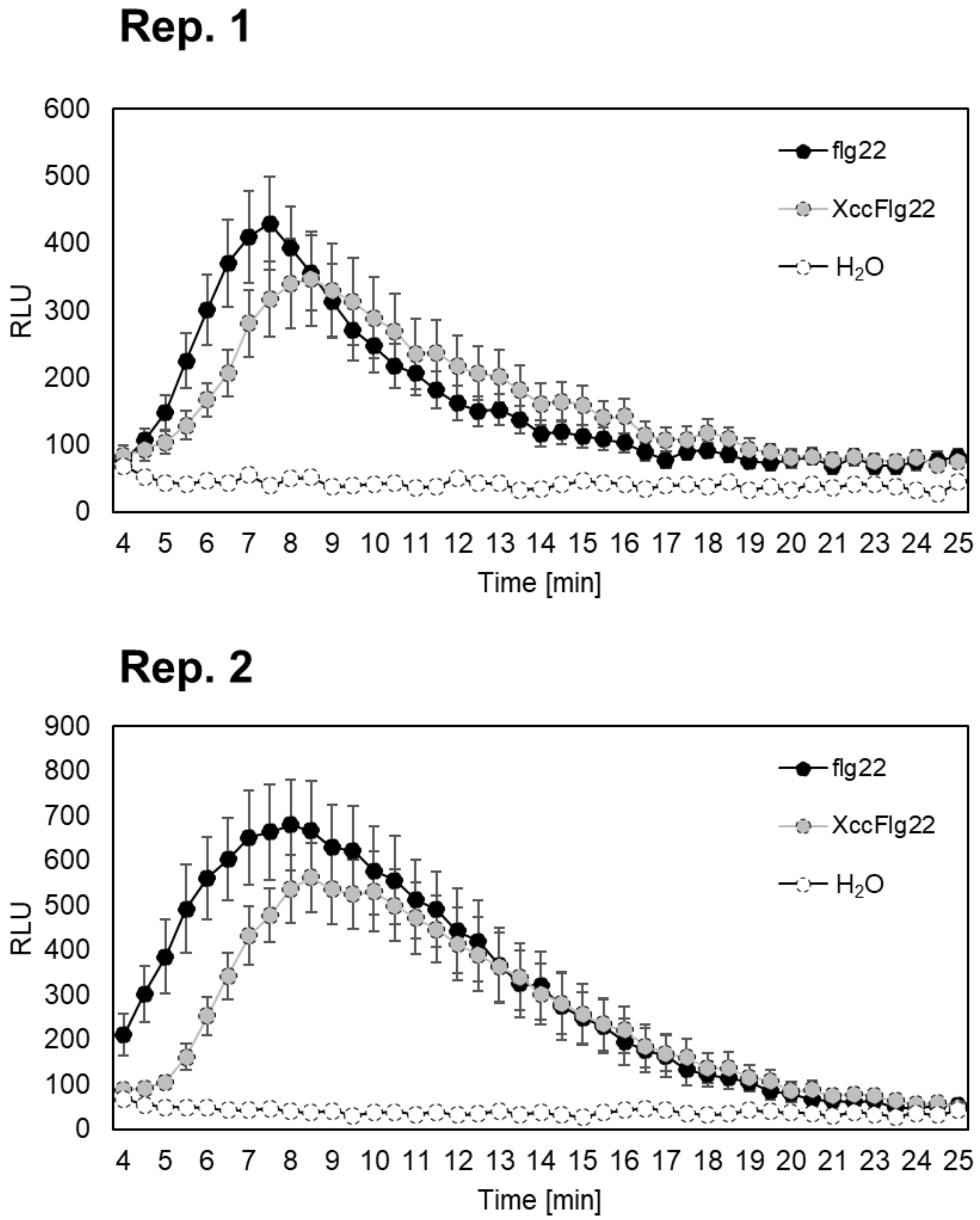


**Figure S9. ROS burst in Gerlach triggered by treatment with distinct flg22.** Discs were cut from 5-6 week old Gerlach and *A. thaliana.* The data represent the means and SEM; n=8-16 discs. The show two biological experiments.

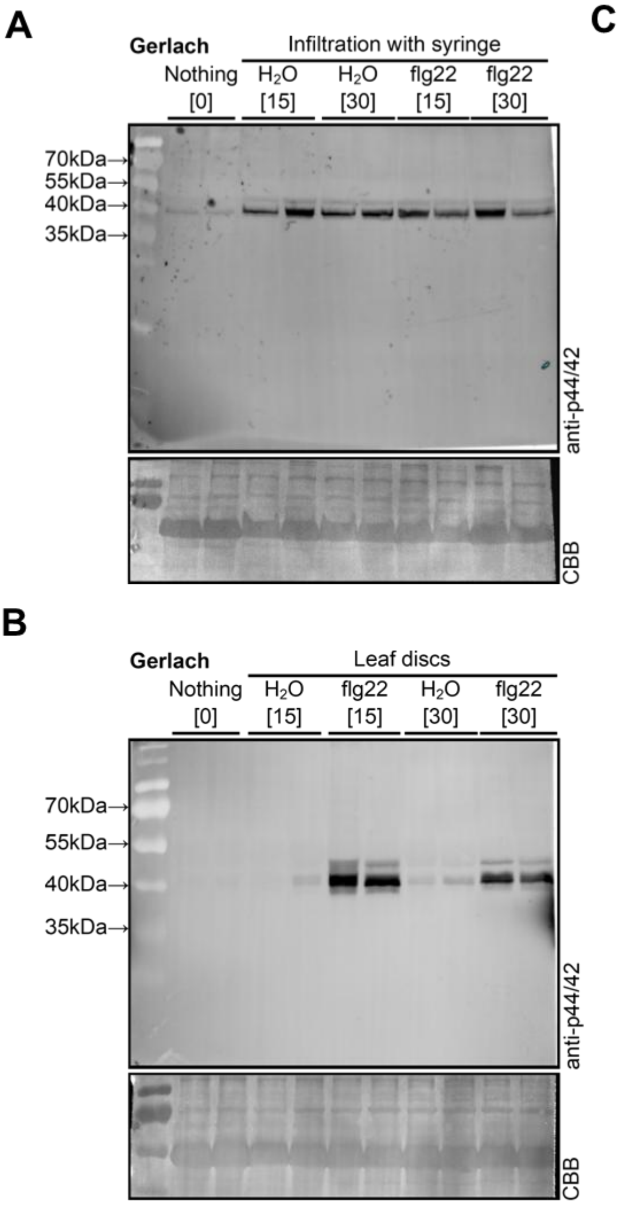


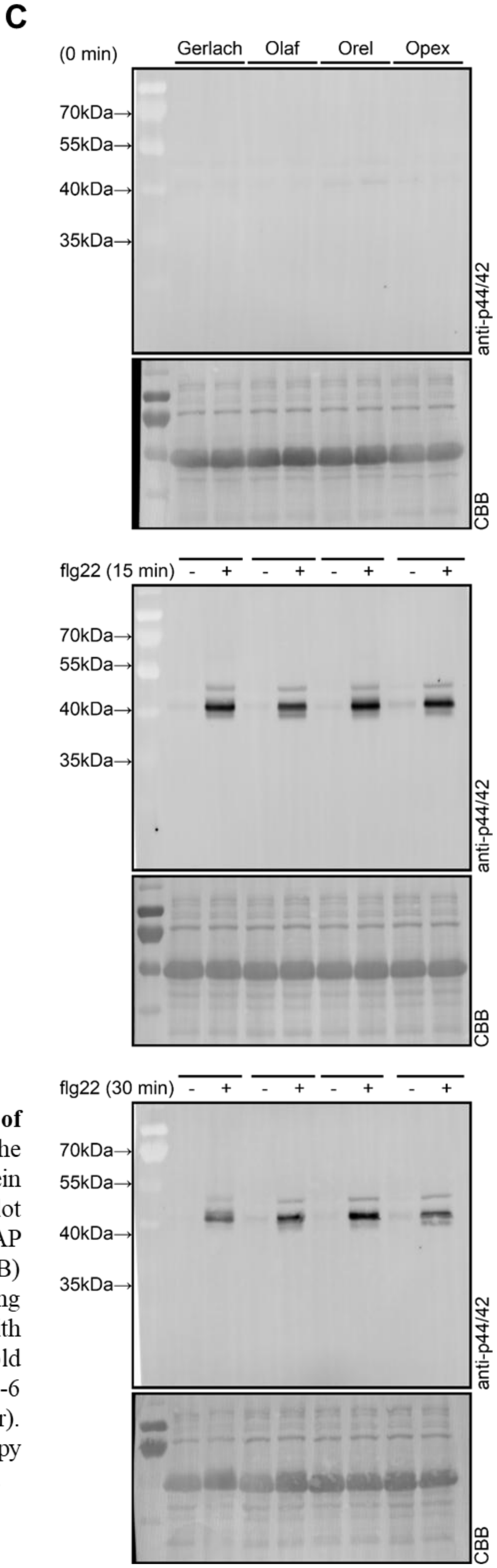


**Figure S10. Original documentation of full gels of immunoblots which were shown in figure 3.** The activation of the putative mitogen-activated protein kinases (MAPKs) was visualized by western blot analysis using the the pERK phospho-p44/42 MAP kinase antibody. Comassie briliant blue (CBB) staining of the membrane served as a loading control. **A)** Needleless syringae infiltration with 5µM flg22 (or water) of leaves from 5-6 week old poppy plants. **B)** Treatment of leaf discs from 5-6 week old Gerlach plants with 5µM flg22 (or water). **C)** Treatment of leaf discs from 5-6 week old poppy plants (four genotypes) with 5µM flg22 (or water).

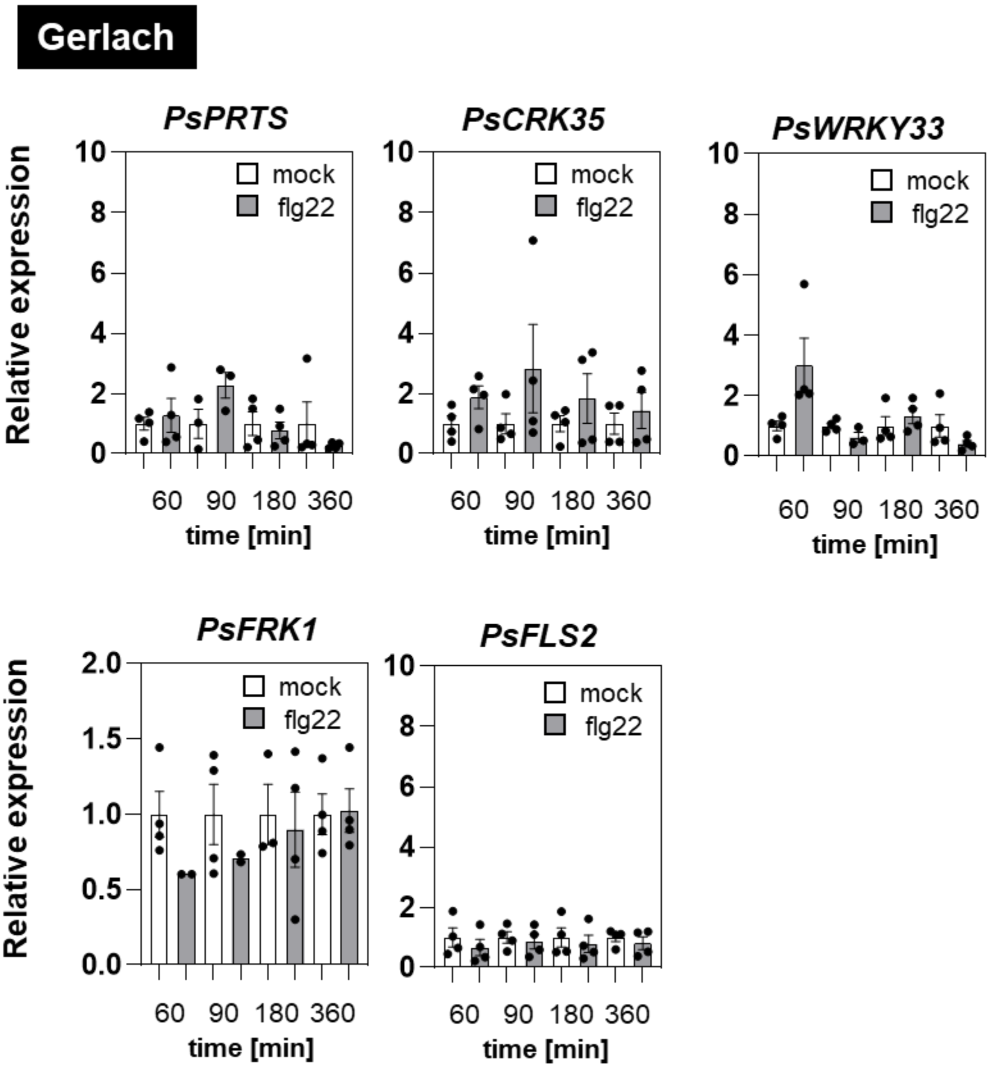


**Figure S11 Gene expression after treatment with flg22.** The leaf discs were cut from 5-6 week old poppy and treated with 5 µM flg22 for 60, 90, 180, 360 minutes. The relative transcription for controls ($H_2O$ treated samples) was normalized to mean 1. The data represent the means + SEM; n=3-4. Mock and flg22 treated samples were tested using two-tailed Student´s t-test. No significant difference ($P<0.05$) was observed.

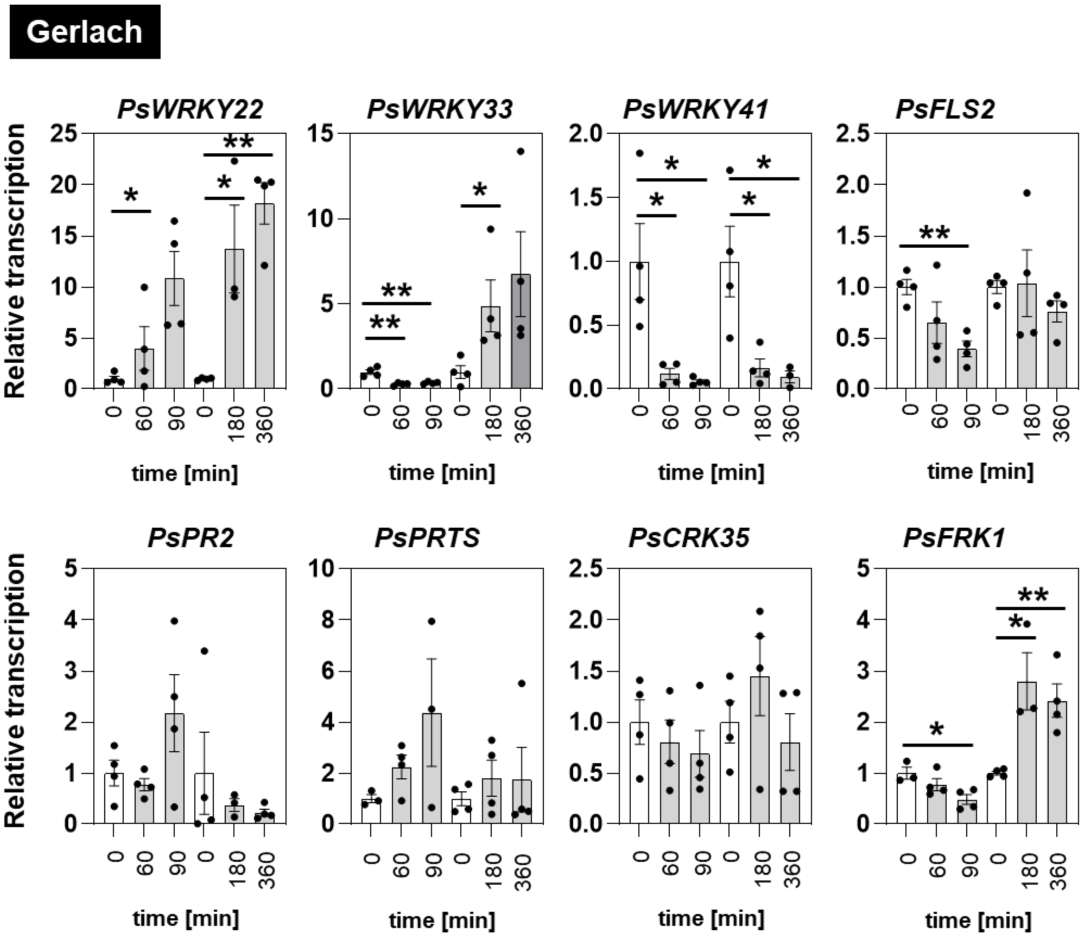


**Figure S12. Gene expression after treatment with $H_2O$ compared to non-treated samples.** The leaf discs were cut from 5-6 week old poppy (cultivar Gerlach) and left overnight in darkness and after that treated with $H_2O$ for 60, 90, 180, 360 minutes. The relative transcription for controls (no treated, directly harvested samples) was normalized to mean 1. The data represent the means + SEM; n=3-4. Asterisks indicate that the mean value is siginificantly different from the control conditions (two-tailed Student´s t-test, *P<0.05, **P<0.01).

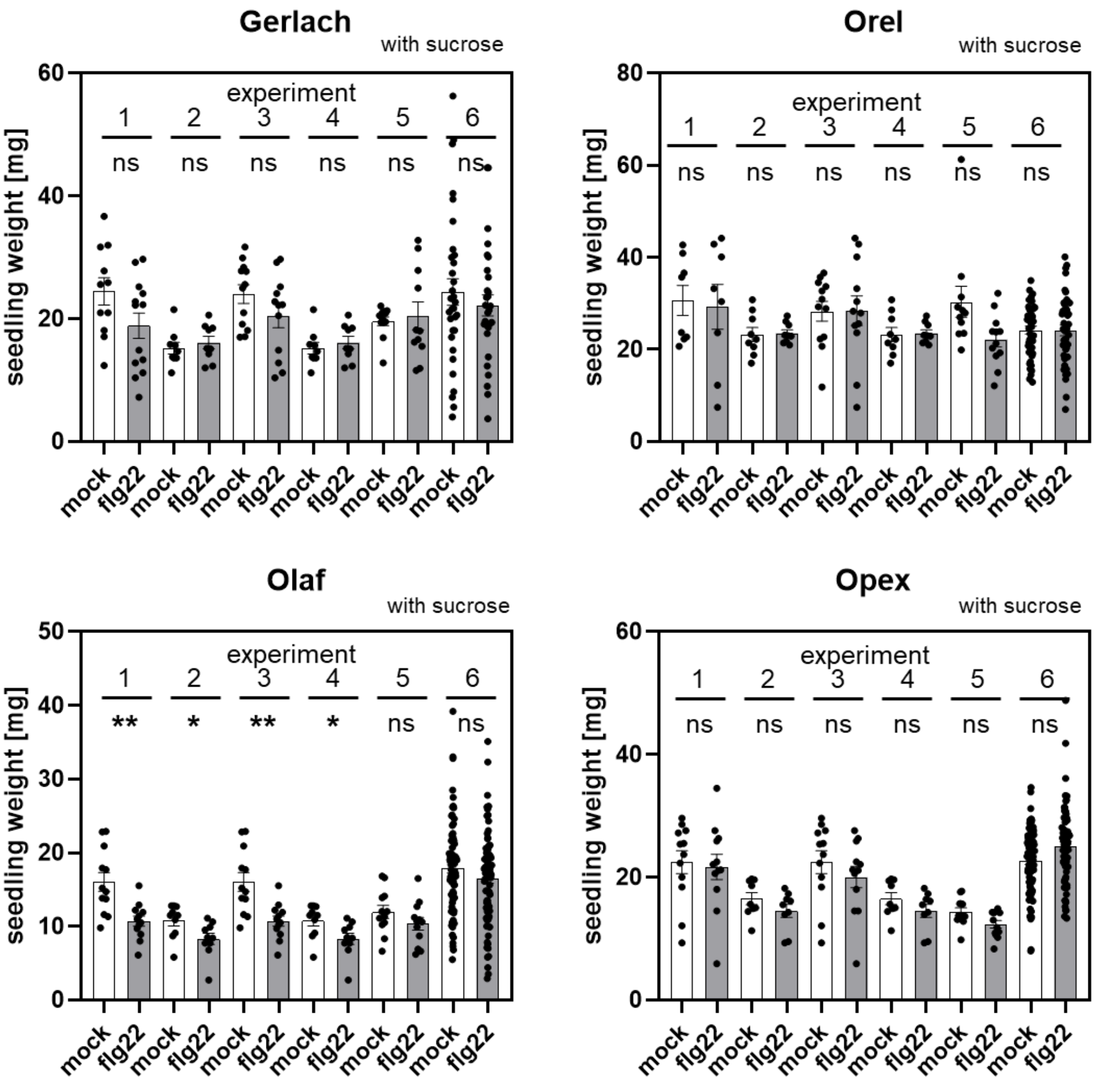


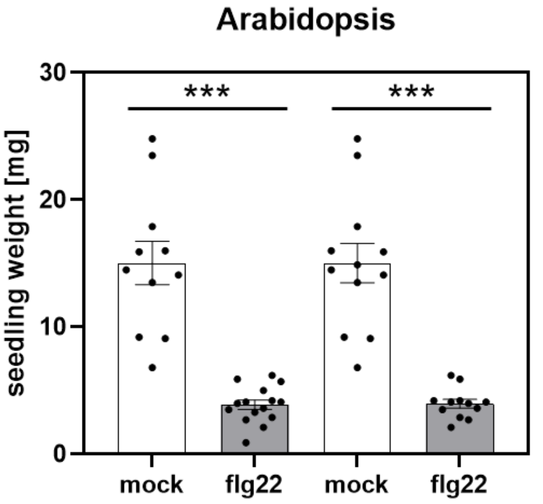


**Figure S13 Raw data for seedlings growth inhibition in media containing sucrose shown in figure 5.** Asterisks indicate that the mean value is siginificantly different from the respektive control conditions (two-tailed Student´s t-test, *P<0.05, **P<0.01, ***P<0.001).

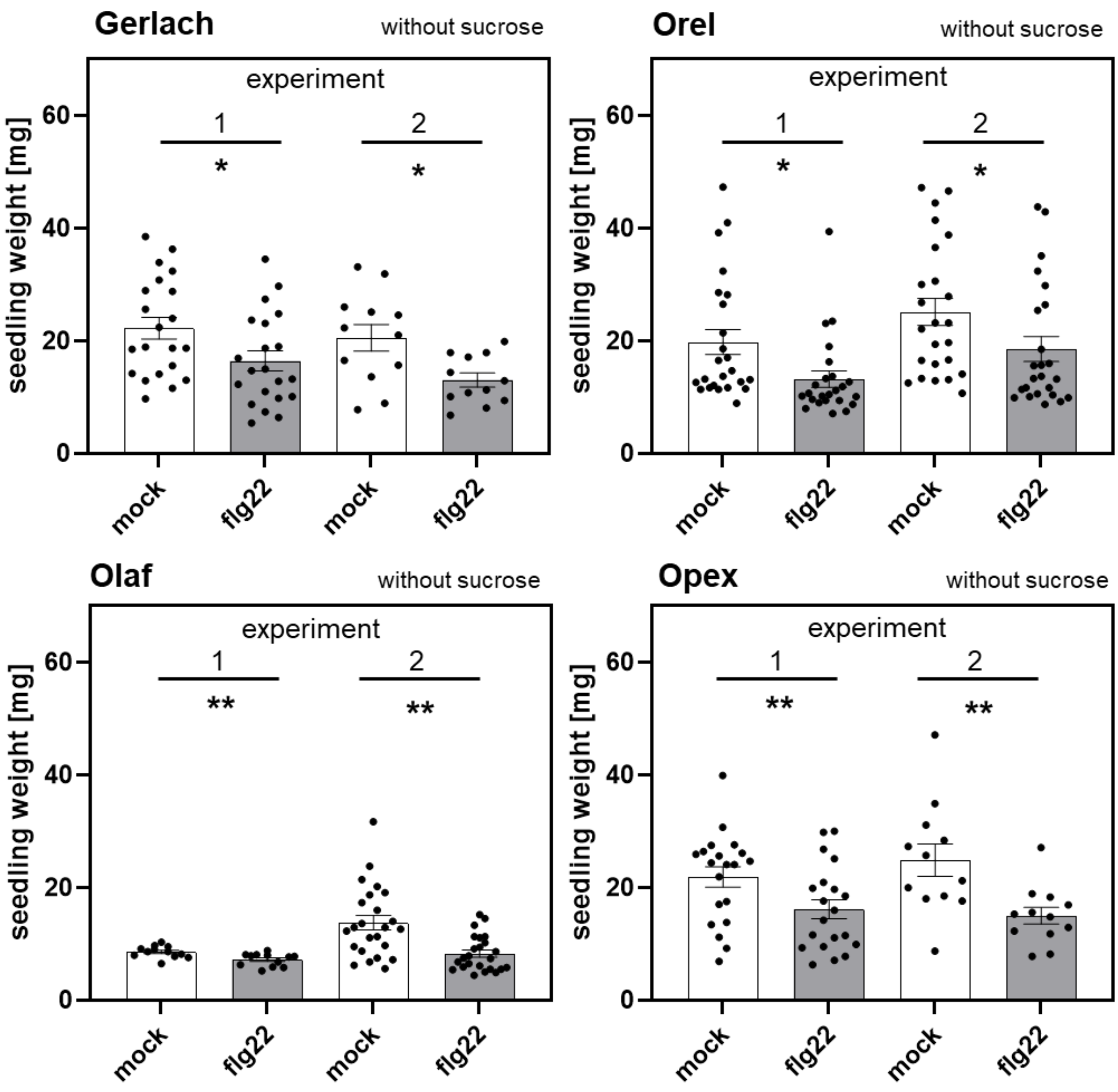


**Figure S14 Raw data for seedlings growth inhibition in media without sucrose shown in figure 5.** Asterisks indicate that the mean value is siginificantly different from the control conditions (two-tailed Student´s t-test, *P<0.05, **P<0.01).

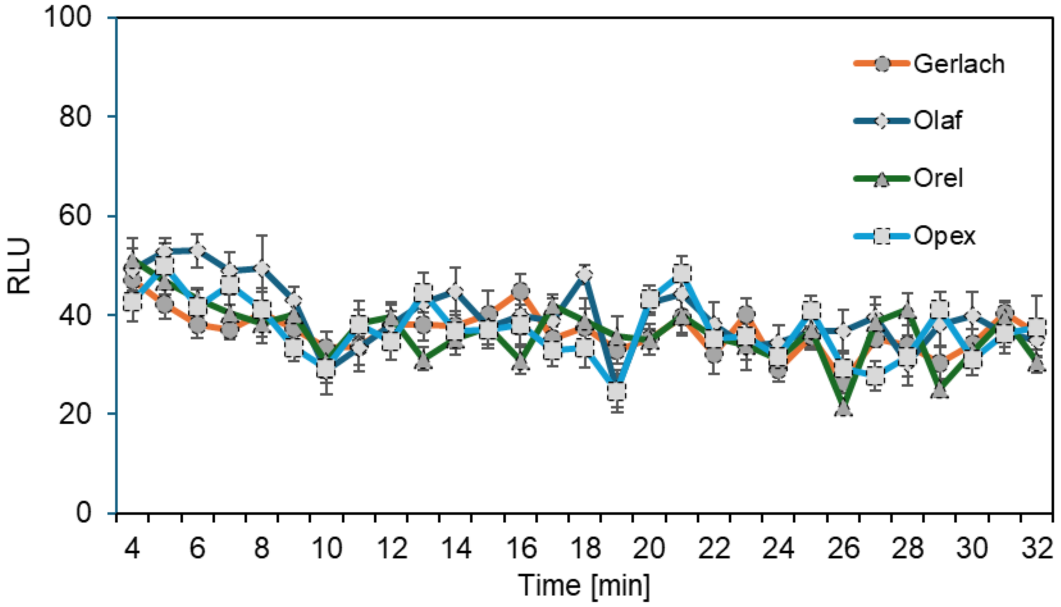


**Figure S15 ROS burst in seedlings of four poppy cultivars after flg22 treatment.** 7-day old seedlings were treated with 5µM flg22. The data represent the means and SEM; n=12 seedlings.

**Table S1. Primers used for RT-qPCR in *Papaver somniferum***

| Accession (NCBI) | (putative) Gene name | Primer name | Orientation | Sequence | Product lenght | Reference | Notes |
|---|---|---|---|---|---|---|---|
| LOC113344176 | *PsWRKY22* | qPCR_PsWRKY22_FP | Forward | TCCAAGAGGATATTACAGATGC | 124 | this study | WRKY transcription factor 22 |
| | | qPCR_PsWRKY22_RP | Reverse | AACGGCATGATTATGTTCAG | | this study | |
| LOC113329236 | *PsWRKY33* | qPCR_PsWRKY33_FP | Forward | CTCTTCCTCCAATATTCTTCCA | 122 | this study | Probable WRKY transcrioption factor 33 |
| | | qPCR_PsWRKY33_RP | Reverse | CTGTTTCTTGCCTGATTGTG | | this study | |
| LOC113321464 | *PsWRKY41* | qPCR_PsWRKY53_FP | Forward | GGGTCTCCGTCGATTCGTTGA | 130 | Ray et al. 2021 | In Ray et. al. named according to soybean as probable WRKY53. |
| | | qPCR_PsWRKY53_RP | Reverse | ACAGCTGGAGAAAGTACGGGC | | Ray et al. 2021 | |
| LOC113355437 | *PsFLS2* | qPCR_PsFLS2_FP | Forward | CTCAACACTGAGCTCCTACG | 113 | this study | LRR receptor-like serine/threonine protein kinase |
| | | qPCR_PsFLS2_RP | Reverse | TCCCTGCATAGTTGTGTCC | | this study | |
| LOC113274805 | *PsPR2* | qPCR_PsPR2_FP | Forward | AAATGTATTCCTTGAAGCGGC | 158 | this study | Putative glucan endo-1,3-beta-glucosidase |
| | | qPCR_PsPR2_RP | Reverse | CTCGTAGGGCATTTAGGGC | | this study | |
| LOC113282088 | *PsPRTS* | qPCR_PsPRTS_FP | Forward | TGACGACCCCAGAATCTCCC | 113 | Ray et al. 2021 | In Ray et al. named according soybean genome as Pathogenesis-related thaumatin superfamily |
| | | qPCR_PsPRTS_RP | Reverse | CAGCCAAGTGCAGGGAAACC | | Ray et al. 2021 | |
| LOC113329942 | *PsCRK35* | qPCR_PsCRK1_FP | Forward | TGGCGAAGGTGGATTTGGAT | 107 | Ray et al. 2021 | In Ray et al. named according soybean genome as Cysteine-rich RLK isoform 1 |
| | | qPCR_PsCRK1_RP | Reverse | ACTCTTGTTCGCCTTGTCCA | | Ray et al. 2021 | |
| LOC113321890 | *PsFRK1* | qPCR_PsFRK1_FP | Forward | CGTGGGTAAACTGTAACTCGG | 135 | this study | Probable LRR receptor-like serine/threonine protein kinase |
| | | qPCR_PsFRK1_RP | Reverse | AATTCCCGCCAAGTCTTAGC | | this study | |
| LOC113323526 | *Actin* | qPCR_Actin_FP | Forward | GCTGTCCTTTCCCTCTACGC | 155 | Zhang et al. 2020 | Actin-11 |
| | | qPCR_Actin_RP | Reverse | AGGGCATCAGTAAGGTCACG | | Zhang et al. 2020 | |

**Table 2 List of binding affinity energies for *P. somniferum* and *A. thaliana* FLS2 and flg22 and *Xcc*Flg22**

| # model | Ligand | Binding Affinity (kcal/mol) | *in silico* Kd (µM) |
|---|---|---|---|
| 0 | *Ps*FLS2+*Xcc*Flg22 | -7.4 | 3.605 |
| 1 | *Ps*FLS2+*Xcc*Flg22 | -7.2 | 5.058 |
| 2 | *Ps*FLS2+*Xcc*Flg22 | -7.2 | 5.058 |
| 3 | *Ps*FLS2+*Xcc*Flg22 | -7.1 | 5.992 |
| 4 | *Ps*FLS2+*Xcc*Flg22 | -6.9 | 8.408 |
| 5 | *Ps*FLS2+*Xcc*Flg22 | -6.9 | 8.408 |
| 6 | *Ps*FLS2+*Xcc*Flg22 | -6.9 | 8.408 |
| 7 | *Ps*FLS2+*Xcc*Flg22 | -6.8 | 9.960 |
| 8 | *Ps*FLS2+*Xcc*Flg22 | -6.8 | 9.960 |
| 0 | *Ps*FLS2+flg22 | -6.5 | 16.555 |
| 1 | *Ps*FLS2+flg22 | -6.3 | 23.230 |
| 2 | *Ps*FLS2+flg22 | -6.2 | 27.517 |
| 3 | *Ps*FLS2+flg22 | -6.1 | 32.595 |
| 4 | *Ps*FLS2+flg22 | -6.1 | 32.595 |
| 5 | *Ps*FLS2+flg22 | -6.1 | 32.595 |
| 6 | *Ps*FLS2+flg22 | -6 | 38.611 |
| 7 | *Ps*FLS2+flg22 | -6 | 38.611 |
| 8 | *Ps*FLS2+flg22 | -6 | 38.611 |
| 0 | *At*FLS2+flg22 | -13.4 | 0.000139 |
| 1 | *At*FLS2+flg22 | -12.9 | 0.000325 |
| 2 | *At*FLS2+flg22 | -12.7 | 0.000456 |
| 3 | *At*FLS2+flg22 | -12.4 | 0.000757 |
| 4 | *At*FLS2+flg22 | -12.4 | 0.000757 |
| 5 | *At*FLS2+flg22 | -12.3 | 0.000897 |
| 6 | *At*FLS2+flg22 | -12 | 0.001491 |
| 7 | *At*FLS2+flg22 | -11.9 | 0.001766 |
| 8 | *At*FLS2+flg22 | -11.9 | 0.001766 |
